# *Pectobacterium sinaloense* sp. nov., a novel phytopathogenic species isolated from potato plants in Mexico

**DOI:** 10.1101/2025.06.20.660644

**Authors:** Jose Luis Valdez-Lopez, Noe Leonardo Palafox-Leal, Glenda Santos-Lopez, Elisa Ines Fantino, Irena Kukavica, Roger C. Levesque, Jesús Méndez-Lozano, Carlos Ignacio Mora-Zamudio, Edgar Antonio Rodríguez-Negrete, Maria Elena Santos-Cervantes, Norma Elena Leyva-Lopez, Edel Pérez-López

**Affiliations:** Instituto Politécnico Nacional, CIIDIR-Sinaloa, Departamento de Biotecnología Agrícola, Guasave, Sinaloa, Mexico; Département de phytologie, Faculté des sciences de l’agriculture et de l’alimentation, Université Laval, Québec City Québec, Canada; Centre de recherche et d’innovation sur les végétaux, Université Laval, Québec City Québec, Canada; Institute de Biologie Intégrative et des Systèmes (IBIS), Université Laval, Québec City Québec, Canada; L’Institute EDS, Université Laval, Quebec City, Québec City, Québec, Canada; Département de microbiologie-infectiologie et d’immunologie, Faculté de médecine, Université Laval, Québec City Québec, Canada; RH PRODUCE, Los Mochis, Sinaloa, Mexico

**Keywords:** Stem rot, Blackleg of potato, Soft rot, *Pectobacterium sinaloense*, Comparative genomics, Potato pathogens, Taxonomic characterization

## Abstract

As part of a broader effort to survey and characterize the diversity of pectolytic bacteria affecting potato crops in Mexico, phytopathogenic strains were isolated from soft rot symptoms in potato plants in Sinaloa. Among them, an atypical *Pectobacterium*-like strain, LFLA-215^T^, could not be confidently assigned to any known species through biochemical or molecular methods. To clarify its taxonomic position and explore its genomic and functional features, whole-genome sequencing and comparative analyses were conducted, accompanied by biochemical, morphological and pathogenicity evaluations. The strain LFLA-215^T^ is Gram-stain-negative, with peritrichous flagella, catalase-positive, and oxidase-negative. Phylogenetic analyses based on the 16S rRNA operon, *dnaJ*, and 923 core genes, confirmed that strain LFLA-215^T^ belongs to the genus *Pectobacterium*. However, genomic similarity values with other *Pectobacterium* species, ranging from 87.73–93.53% (ANIb), 87.63–93.46% (ANIu), and 34.0–52.1% (isDDH), fell below species delineation thresholds. *Pectobacterium colocasium* LJ1^T^ showed the closest relationship to LFLA-215^T^, whereas *Pectobacterium parmentieri* RNS 08-42-1A^T^ was the most distantly related. Although LFLA-215^T^ fulfilled Koch’s postulates and demonstrated pathogenicity in potato plants, its virulence on tubers was comparatively lower than that of other known *Pectobacterium* strains, which could be related to the size and the reduction of the total number of genes when analyzed its complete genome reported here. Taken all together, our findings support the classification of strain LFLA-215^T^ as a novel species within the genus *Pectobacterium*, for which the name *Pectobacterium sinaloense* sp. nov. is proposed, with LFLA-215^T^ designated as the type strain.

## INTRODUCTION

Over the last decade, soft-rot causing *Pectobacterium* species have ranked among the top ten most studied plant pathogenic bacteria (Mansfield et al., 2012). With a broad host range, affecting nearly 35% of angiosperms, these bacterial phytopathogens are responsible for soft rot diseases in many economically important crops, leading to estimated annual losses of $1 billion and threatening food security and the global economy (Domingo et al., 2021; Marquez-Villavicencio et al., 2011). In potato (*Solanum tuberosum* L.), they cause soft rot, blackleg, wilt, and aerial stem rot (Toth et al., 2021). Their pathogenicity relies on a diverse arsenal of plant cell wall-degrading enzymes (PCWDEs) such as pectinases, cellulases, proteases, which enable tissue maceration (Oulghazi et al., 2019; Waleron et al., 2019). Although *Pectobacterium* is not considered a stealth pathogen, it can benefit from stealth-like strategies, including the induction of host susceptibility responses and the formation of heterogeneous populations with specialized roles (Gorshkov & Parfirova, 2023; Panda et al., 2016). Disease development is further promoted by environmental conditions such as high humidity, elevated temperatures, and hypoxia due to waterlogged soils (Maciag et al., 2024).

Recently, the taxonomy of *Pectobacterium* has undergone extensive revisions, driven by advances in phylogenomic analyses (Oulghazi et al., 2019; Pasanen et al., 2020; Portier et al., 2019, 2020; Sawada et al., 2024). As of March 2025, the *List of Prokaryotic Names with Standing in Nomenclature* (LPSN) recognizes 22 validly published *Pectobacterium* species (Parte et al., 2020). Among them, *P. cacticida* has been reclassified under a new genus as *Alcorniella cacticida* comb. nov. (Jonca et al., 2024), while *P. carnegieanum* is considered likely to fall out of use due to the lack of a valid type strain (Young, 2011). Despite these developments, *Pectobacterium* remains a dynamic taxonomic group, with ongoing efforts to resolve its diversity and evolutionary relationships.

In Mexico, potato cultivation is a key pillar of national agriculture, with an annual production value nearing one billion dollars. The state of Sinaloa alone accounts for over 21.5% of the country’s total production (SIAP, 2024). However, *Pectobacterium* species remain a persistent threat in this region, causing significant damage to potato crops (Palafox-Leal et al., 2024; Santos-Cervantes et al., 2024; Valdez-Lopez et al., 2025). As part of a broader effort to survey and characterize the diversity of pectolytic bacteria affecting potato in Mexico, several *Pectobacterium* strains were isolated. Preliminary phenotypic and genotypic analyses identified one atypical strain, LFLA-215^T^, with reduced virulence that could not be assigned to any currently known *Pectobacterium* species. This strain was isolated in January-2020 from a potato stem sample showing blackleg symptoms collected in Ahome, Sinaloa, Mexico (25.8656 N, 108.9249 W). To clarify its taxonomic status and explore its genomic and functional traits, we conducted whole-genome sequencing, comparative genomics, and phenotypic characterization including biochemical, morphological, pathogenicity, and virulence assays. Our results support the designation of this isolate as a novel species, for which we propose the name *Pectobacterium sinaloense* sp. nov.

## GENOMIC FEATURES OF STRAIN LFLA-215^T^

For whole-genome sequencing, strain LFLA-215^T^ was grown in Luria-Bertani (LB) broth at 28 °C for 18 h. Genomic DNA was extracted using the lysis buffer described by Chen and Kuo (1993), followed by alcohol precipitation. A hybrid sequencing approach combining Oxford Nanopore Technologies (ONT) and Illumina platforms was employed. ONT libraries were prepared using the v14 chemistry kit, and raw reads (2.1 Gbp) were filtered with Filtlong v0.2.1 (default parameters; https://github.com/rrwick/Filtlong), resulting in a high-quality read (HQR) subset of 457 Mbp. Assembly was performed using Flye v2.9.1 (Kolmogorov et al., 2019) with the HQR subset and parameters optimized for long read data. The resulting assembly was polished with Medaka v1.8.0 (Oxford Nanopore Technologies Ltd., 2018) using the same HQR subset. Illumina reads over 2.4 Gbp, were then used for additional polishing with Polypolish v0.6.0 (Bouras et al., 2024).

Annotation of the LFLA-215⍰ genome was conducted using the Nextflow Mettannotator Pipeline v1.3, a comprehensive framework that integrates multiple tools and databases for functional and structural genome characterization (**Table S1**) (Gurbich et al., 2025). The pipeline was executed with the *-bakta* parameter. Genome completeness was assessed using the *Pectobacterium*-specific dataset integrated into CheckM v1.2.3 (Parks et al., 2015). Secretion system types I, II, III, IV, and VI were predicted using a pipeline developed for Gram-negative bacteria, which incorporates MacSyFinder v2 (Zhang et al., 2023; Néron et al., 2023). In addition, a manual curation of the merged annotation output from Mettannotator v1.3 was performed to identify genes encoding plant cell wall-degrading enzymes (PCWDEs) and potential toxins.

The genome of strain LFLA-215^T^ (**Figure 1A-B, Table S2**) was assembled at the chromosome level, with a final coverage of 632x, and no plasmids detected. It spans 4,522,015 bp with a G+C content of 51.6 %. A total of 4,043 protein-coding sequences were annotated, resulting in a coding density of 0.86; among these, 164 were classified as hypothetical proteins. Additionally, 77 tRNAs and 22 rRNAs were identified. Genome quality assessment using CheckM indicated 93.33% completeness and 0.44% contamination. The relatively low completeness score likely reflects the limited representation of the *Pectobacterium* genomes in the CheckM reference dataset, which includes only five genomes.

**Figure 1.**
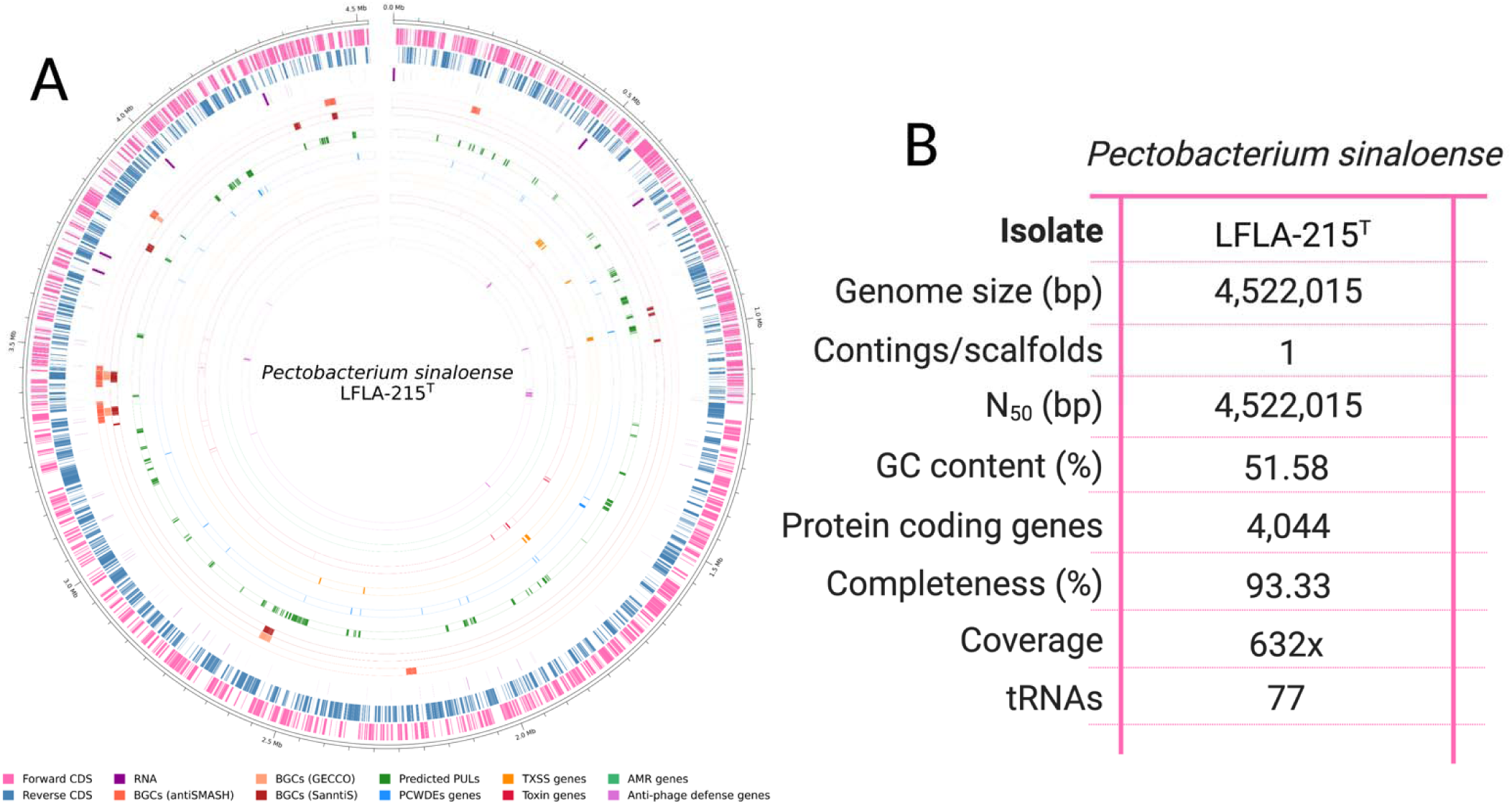
Chromosomal architecture and functional annotation of *Pectobacterium sinaloense* LFLA-215. (A) Circos plot generated with pyCirclize 1.4.0 (Shimoyama, Y., 2022), depicting chromosomal features, arranged from outer to inner rings: sequence coordinates, coding sequences (CDS) on positive/negative strands, RNA features, biosynthetic gene clusters (BGCs) predicted by antiSMASH, GECCO, and SanntiS, polysaccharide utilization loci (PULs), plant cell wall-degrading enzymes (PCWDEs) genes, type I, II, III, and VI secretion systems (TXSS), toxin genes, antimicrobial resistance (AMR) determinants and anti-phage defense systems. (B) Summary table of genome assembly metrics, completeness (using CheckM), and annotation statistics.

Key genomic features of LFLA-215⍰ include biosynthetic gene clusters (BGCs) and polysaccharide utilization loci (PULs) (**Table S3**). BGCs were detected using antiSMASH (Blin et al., 2023), Gecco (Carroll et al., 2021), and SanntiS (Sanchez et al., 2023), revealing clusters involved in the biosynthesis of β-lactone-containing protease inhibitors, thiopeptides, non-ribosomal peptide synthetase (NRPS)-Type I polyketide synthase (T1PKS) hybrids, NRPS-metallophores, and NRPS-independent IucA/IucC-like siderophores (NI-siderophores) (**Table S3**). These clusters likely provide competitive advantages by enabling the production of bioactive molecules such as antibiotics, toxins, or siderophores which support microbial competition and niche establishment. A genomic region associated with the homoserine lactone production was also identified. These molecules function as *quorum sensing* (QS) signals, regulating the expression of genes linked to virulence, biofilm formation, and coordinated behavior. In *Pectobacterium*, QS has been shown to control over 70 regulatory elements, including those governing PCWDE production and type I, II, III, and VI secretion systems (Liu et al., 2008; Van Gijsegem et al., 2021).

PULs were predicted using dbCAN3 (Zheng et al., 2023), revealing 63 loci associated with the degradation of various carbohydrates. Substrate-specific PULs included those targeting pectin (10), cellobiose (4), levoglucosan (3), xylan (3), host glycans (2), galactan (2), starch (2), and one locus each for arabinan, β-glucoside, capsular polysaccharide, cellulose, chitin, glycogen, glycosaminoglycan, and melibiose and 29 with no description available (**Table S3**). In parallel, 33 coding sequences corresponding to PCWDEs were identified, including 27 pectinases, two cellulases, and four proteases (**Table S4**). Notably, most of these genes were co-localized within PULs predicted by dbCAN3 (**Figure 1A**), suggesting that PUL prediction tools can be leveraged to discover novel PCWDEs based on substrate specificity and conserved catalytic domains. The dominance of pectinases in the PCWDEs repertoire aligns with previous findings that *Pectobacterium* species harbor an expanded set of pectin-degrading enzymes compared to other soft rot pathogens (Jonkheer et al., 2021), a feature likely contributing to the genus’s aggressive maceration phenotype.

An important pathogenicity-related feature of *Pectobacterium* species is the presence of multiple secretion systems and associated toxins. In strain LFLA-215⍰, six secretion systems were identified (**Table S4**): three type I (T1SS), and one each of type II (T2SS), type III (T3SS), and type VI (T6SS). No type IV secretion system (T4SS) was detected through either automated or manual screening. Secretion systems are considered the second most critical virulence determinants in *Pectobacterium*, after PCWDEs (Arizala & Arif, 2019; Van Gijsegem et al., 2021). In total, 34 toxin-related genes were identified (**Table S4**), including the well-characterized necrosis-inducing protein *Nip*, a conserved protein that promotes plant cell death and facilitates disease progression (Charkowski et al., 2012). These elements, PCWDEs, secretion systems (TXSS), and toxins, form the core arsenal driving soft rot pathogenicity in *Pectobacterium* (Xu et al., 2021).

Beyond virulence, other functional categories were annotated. AMRFinderPlus (Feldgarden et al., 2021) identified three genes linked to streptogramin, acid, and arsenate resistance, which may offer baseline protection against environmental stressors. Additionally, DefenseFinder (Tesson et al., 2022) detected 38 genes spanning 11 distinct prokaryotic antiviral defense systems (**Table S3**). The presence of these antiviral defense systems suggests LFLA-215⍰ is equipped to withstand phage infection, likely contributing to its genomic stability in competitive ecological niches.

## TAXONOMIC PLACEMENT OF STRAIN LFLA-215^T^

For preliminary taxonomic identification, the complete 16S rRNA gene sequence of strain LFLA-215⍰ (GenBank accession number: PV590475.1) was extracted from the genome annotation file and queried against the NCBI RefSeq genome database using BLASTn. The highest similarity was found within the *Pectobacterium* genus, showing 100% identity and 100% coverage with *Pectobacterium* sp. strain CFBP8739 (accession number: GCF_013449375.1), a strain that currently lacks species-level classification. Additional high-scoring hits included *Pectobacterium* strains CSR2 and CSR3 (GenBank accession numbers: GCF_048593165.1 and GCF_048593175.1), both annotated as *Pectobacterium colocasium*, showing 99.55% identity with strain LFLA-215⍰. All other matches exhibited lower identity values. Given these results, and the lack of peer-reviewed publications validating strains CFBP8739, CSR2, and CSR3, additional analyses were conducted to further clarify the taxonomic placement of strain LFLA-215⍰ within the *Pectobacterium* genus.

Genome-scale taxonomic comparisons of strain LFLA-215^T^ were performed using *in silico* DNA-DNA hybridization (*is*DDH) via the Type Strain Genome Server (TYGS; https://tygs.dsmz.de), applying the recommended distance formula d4 (Meier-Kolthoff & Göker, 2019). In addition, average nucleotide identity (ANI) was calculated using both OrthoANI USEARCH (ANIu) and OrthoANI BLAST-based (ANIb) algorithms (Lee et al., 2016; Yoon et al., 2017). These three methods, *is*DDH, ANIb, and ANIu, were used to compare LFLA-215^T^ with strains CFBP8739, CSR2, CSR3, as well as with the type strains of all validly described *Pectobacterium* species, and representative species from other genera within the *Pectobacteriaceae* family (**Table 1**). *Budvicia aquatica* strain FDAARGOS_387^T^ was included as an outgroup (**Table 1**).

**Table 1.**
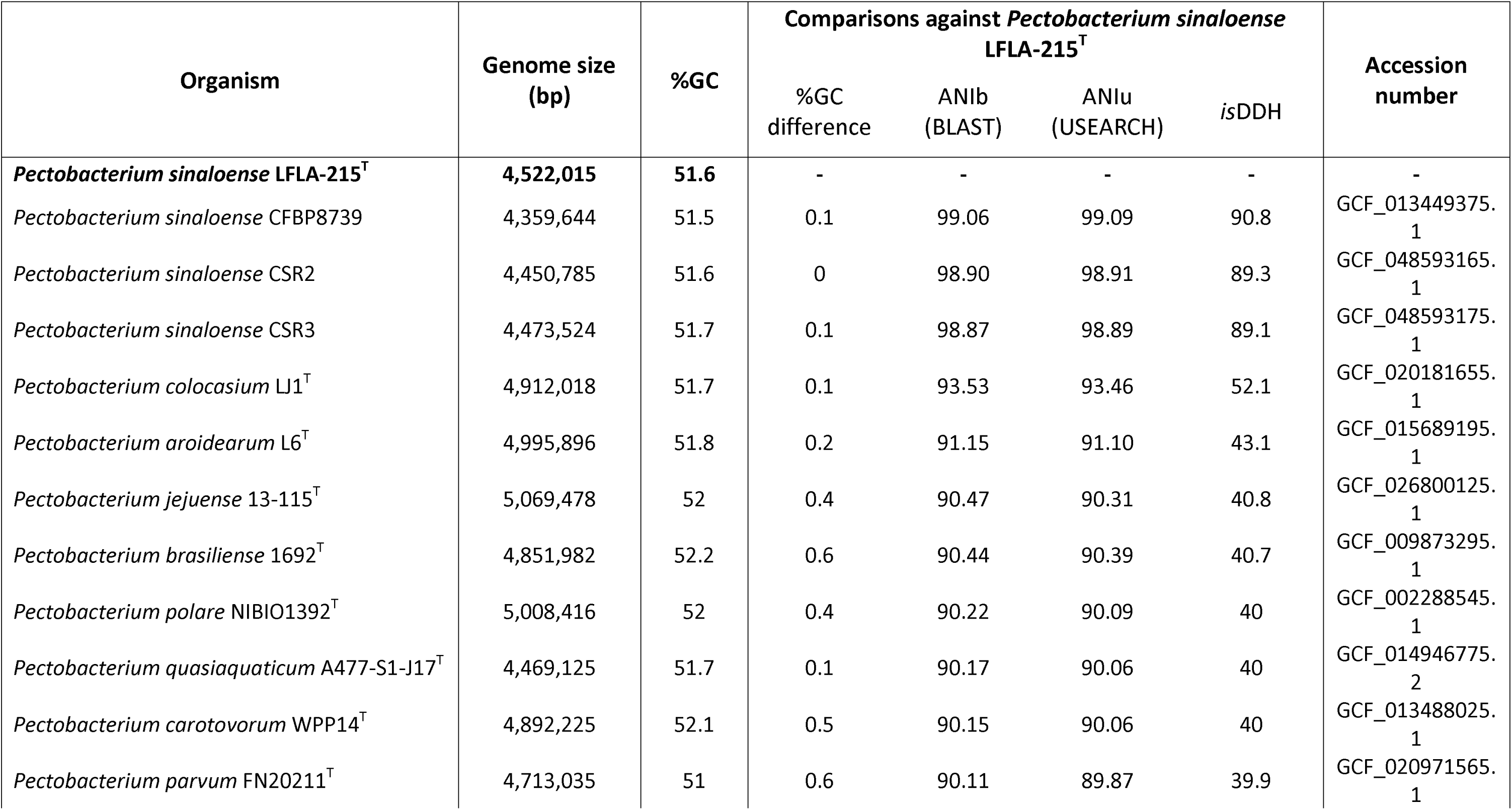
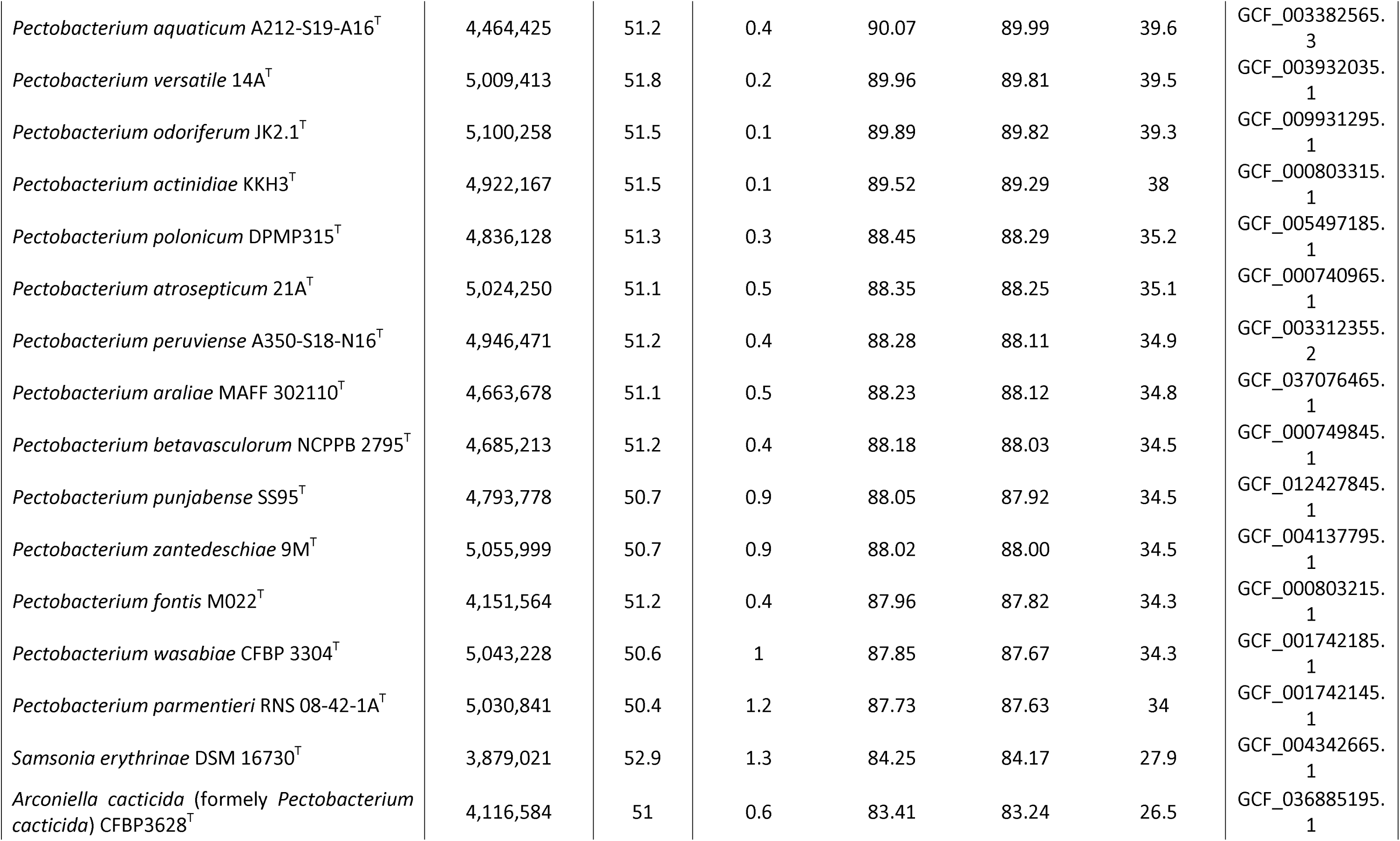
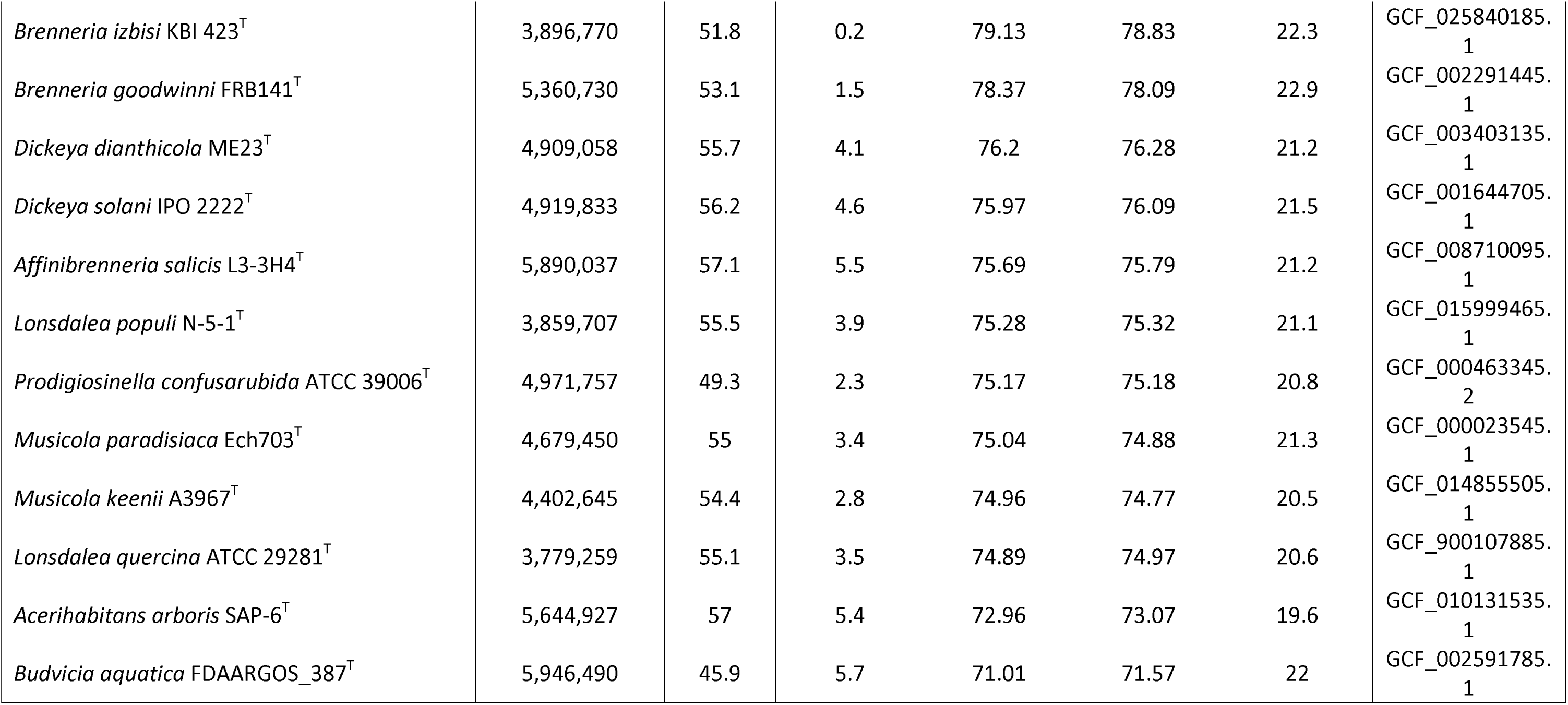
Genomic relationship between strain Pectobacterium sinaloense LFLA-215^T^ and closely related strains.

The results showed that LFLA-215^T^ exhibited ANIb, ANIu, and *is*DDH values of 99.06-98.87%, 99.09-98.89%, and 90.8-89.1%, respectively, when compared with CFBP8739, CSR2 and CSR3, well above the accepted species delineation thresholds of 95-96% for ANI and 70% for *is*DDH (**Table 1**) (Chun et al., 2018). None of the other tested species within the *Pectobacteriaceae* exceeded these thresholds. Among them, *Pectobacterium* species displayed the highest similarity values, ranging from 87.73-93.53% for ANIb, 87.63-93.46% for ANIu, and 34.0-52.1% for *is*DDH (**Table 1**). *Pectobacterium colocasium* LJ1^T^ was the closest relative to LFLA-215^T^, while *Pectobacterium parmentieri* RNS 08-42-1A^T^ was the most distantly related (**Table 1**). These findings support the classification of LFLA-215^T^, along CFBP8739, CSR2 and CSR3, as members of a novel species within the *Pectobacterium* genus. We propose the name *Pectobacterium sinaloense* sp. nov., in reference to the agriculturally productive region of Sinaloa, Mexico, where strain LFLA-215⍰ was isolated.

To further clarify the phylogenetic position of this novel taxon, multiple analyses were conducted using distinct molecular markers. Sequences for the 16S rRNA operon and *dnaJ* gene were aligned using ClustalW, as implemented in MEGA 11 (Tamura et al., 2021), incorporating sequences from 40 strains (representing 37 species) (**Table 1**). In addition, a core genome alignment was generated using Roary (Page et al., 2015), applying a 70% protein identity threshold and requiring 100% gene presence across all genomes. Prior to this step, all 40 genomes were *de novo* re-annotated using Bakta v1.9.4 (Deposited in Zenodo) (Schwengers et al., 2021). Phylogenetic trees based on 16S rRNA, *dnaJ*, and core genome alignments were inferred using IQ-TREE v3.0.0. Model selection was conducted with ModelFinder, and maximum-likelihood trees were generated with 1,000 ultrafast bootstrap replicates (Kalyaanamoorthy et al., 2017; Hoang et al., 2018; Wong et al., 2025). The phylogenetic analyses confirmed the taxonomic status of *P*. *sinaloense* sp. nov. within the *Pectobacteriaceae* family (Figs. 2, S1, S2). In all three analyses, strain LFLA-215^T^ consistently clustered with CFBP8739, CSR2, and CSR3, positioning the proposed species firmly within the *Pectobacterium* genus. Furthermore, the *P. sinaloense* strains consistently formed a heterogeneous clade with *P. colocasium* LJ1^T^ and *P. aroidearum* L6^T^, suggesting these taxa share a common ancestor. The core genome analysis, based on 926 core genes and encompassing 989,863 nucleotide sites, 41.02% of which were parsimony-informative, grouped all *Pectobacterium* type strains into a single clade with 100% ultrafast bootstrap support (**Figure 2**). Similarly, the *dnaJ* gene analysis (**Figure S1**) yielded a consistent clustering pattern, supported by a 97% bootstrap value. In contrast, the 16S rRNA-based analysis (**Figure S2**) failed to clearly resolve the *Pectobacterium* clade, showing overlap with the genus *Alcorniella* (formerly *Pectobacterium* spp.) (Jonca et al., 2024), and with *S. erythrinae* DSM 16730^T^, both representing the closest relatives to *Pectobacterium*. Additionally, bootstrap support values in the 16S rRNA tree were notably lower than those in the *dnaJ* and core-gene trees. These findings align with the observations of Mainello-Land et al. (2024), who demonstrated that the *dnaJ* gene has nearly six times the discriminatory power of the 16S rRNA operon when resolving relationships within the *Pectobacteriaceae* genera *Dickeya* and *Pectobacterium.*

**Figure 2.**
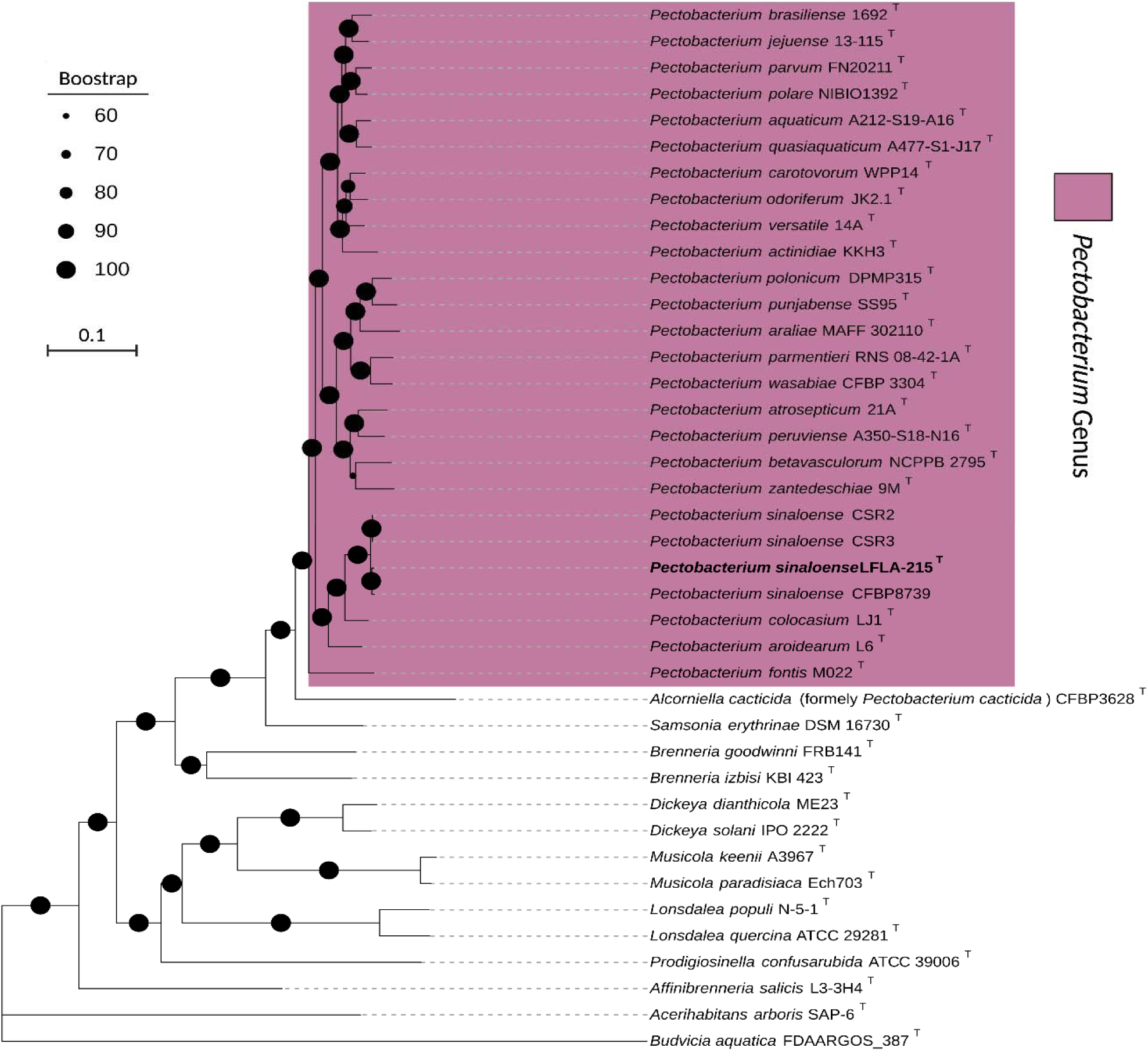
Maximum-likelihood phylogenomic tree reconstructed based on the concatenated alignment of 926 core genes, showing the relationships between *Pectobacterium sinaloense* LFLA-215 (bold type) and closely related strains from the family *Pectobacteriaceae* (listed in **Table 1**). The total alignment length was 989,863 bp and included 40 sequences, with 41.02% parsimony-informative sites. Phylogenetic inference was performed using IQ-TREE v3.0.0 under the GTR+F+I+R7 model with 1,000 ultrafast bootstrap replicates. Ultrafast bootstrap values are indicated at the nodes (only values ≥60 are shown). *Pectobacterium* clade is highlighted in magenta. *Budvicia aquatica* FDAARGOS_387 was used as an outgroup.

## PHENOTYPIC CHARACTERIZATION OF STRAIN LFLA-215^T^

To evaluate the phenotypic traits of strain LFLA-215^T^, a series of routine tests commonly used for *Pectobacterium* species were conducted (Palafox et al., 2024; Silva et al., 2020). These included Gram staining, catalase and oxidase activity, growth at 28⍰°C and 37⍰°C, fluorescence production on B-King medium, and pectolytic activity on potato slices. Motility was assessed using the protocol described by Shields and Cathcart (2011), while osmotic tolerance was tested by culturing the strain in LB broth supplemented with NaCl at concentrations of 1%, 4%, and 8%, and adjusted to pH values of 5, 6, 7, and 8.

Colony and cellular morphology were examined by streaking bacteria onto LB agar and incubating at 28⍰°C; colony images were captured after 24 h. Ultrastructural analysis was performed using transmission electron microscopy (TEM) with a JEOL 2100Plus (Jeol, Japan) at 200 kV and 10,000× magnification. Carbon source utilization and chemical sensitivity were evaluated using BIOLOG Phenotypic Microarray (PM) plates 1, 2, 11C, 12B, 15B, 16A, 17A, 18C, and 20B, following the manufacturer’s instructions. Kinetic responses were recorded over a 72-h period in a BIOLOG OmniLog reader (Biolog, USA), and analyzed using BIOLOG Data Analysis Software 1.7. Assays were performed in duplicate (n = 2 for carbon source assays; n = 8 for chemical sensitivity), and only consistent, reproducible results were kept. Comparative phenotypic data from *Pectobacterium* species, including the most closely related species (*P. aroidearum* and *P. colocasium*), were retrieved from Zhou et al. (2022) and Hong et al. (2023). Pathogenicity and virulence were evaluated on plants and potato tubers, respectively. The strain LFLOG-78 of *Pectobacterium versatile,* previously characterized by us, was included as control (Valdez-Lopez et al., 2025). Pathogenicity assays were performed as previously described (Valdez-Lopez et al., 2025). Bacterial suspensions (10⍰mM MgSO_4_, OD₆₀₀ = 0.8, approximately 10⁸⍰CFU/mL) were injected into the stems of 5-week-old potato plants (cv. Fianna). Each treatment included three plants, along with negative controls inoculated with sterile 10⍰mM MgSO₄. Plants were maintained at 28⍰°C with 80% relative humidity, and symptoms were monitored daily for one week. Koch’s postulates were fulfilled by re-isolating and identifying the inoculated bacteria.

For virulence assays, bacterial suspensions preparation, tuber inoculation, and incubation were carried out as previously described (Han et al., 2023). Cultures were incubated for 18⍰h, centrifugated, washed with 10⍰mM MgSO_4_, and resuspended in the same buffer to an OD_600_ of 0.27. Potato tubers (cv. Fianna) were punctured with a 1000⍰µL pipette tip and 50⍰µL of the bacterial suspension was applied to each wound. Eight tubers were used per isolate, and five were mock-inoculated with buffer as controls. Wounds were sealed with vaseline, and tubers were wrapped in sterile, moistened paper and incubated in plastic containers at room temperature (23–25⍰°C) in darkness for 72⍰h. Tubers were then sliced, and the macerated tissue was scraped and weighed. Control values were used to normalize decay in treated samples. The experiment was repeated twice, and statistical significance was assessed using the Kruskal–Wallis test followed by Dunn’s post hoc test with Bonferroni correction.

After 24⍰h of incubation on LB agar at 28⍰°C, strain LFLA-215^T^ formed small (1–2⍰mm diameter), circular, opaque colonies with a white-beige coloration, entire margins, convex elevation, and a moist surface. No diffusible pigment production was observed (**Figure 3A**). Strain LFLA-215^T^ was also characterized as a Gram-negative, rod-shaped bacterium measuring 4.3 ± 0.9 µm in length and 1.31 ± 0.09 µm in width, with peritrichous flagella (**Figure 3B**, **Table S5**). Motility, catalase activity, and pectolytic activity on potato slices were confirmed, whereas oxidase activity and fluorescence on B-king medium were not detected. The strain grew well in LB broth at both 28⍰°C and 37⍰°C, as well as in media supplemented with 1% NaCl and at pH values ranging from 4 to 8. No growth was observed at 4% or 8% NaCl.

**Figure 3.**
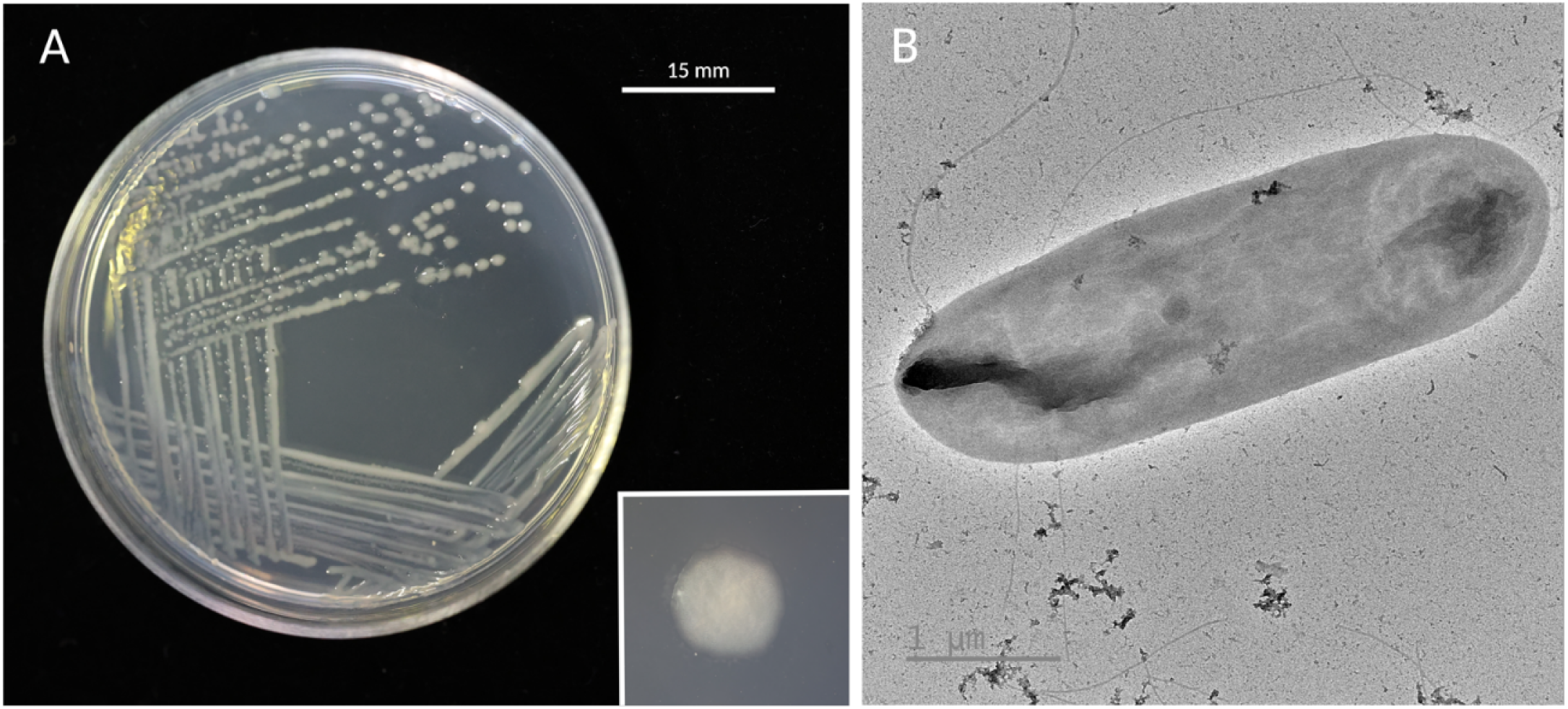
Colony and cellular morphology of *Pectobacterium sinaloense* LFLA-215. (A) Colony morphology on LB agar after incubation at 28⍰°C for 24⍰h. Colonies are small (1–2⍰mm in diameter), circular, opaque, and white-beige in color, with entire margins, convex elevation, moist surfaces, and no diffusible pigment production. (B) Transmission electron micrograph, showing a rod-shaped cell with peritrichous flagella.

Strain LFLA-215⍰ tested positive for the utilization of a broad spectrum of carbon sources, including various sugars, amino acids, sugar alcohols, organic acids, cyclodextrins, and complex polysaccharides such as pectin, inulin, glycogen, and laminarin. This extensive metabolic versatility includes substrates like L-arabinose, D-galactose, D-glucuronic acid, D-mannose, glycerol, L-proline, sucrose, maltose, and N-acetyl-D-glucosamine, among others (**Tables 2, S6**). The strain tested negative for the utilization of propionic acid, capric acid, and D-serine. Weak positive reactions were observed for α-ketobutyric acid, 2-hydroxybenzoic acid, itaconic acid, L-methionine, and 2,3-butanedione (**Tables 2, S6**).

In chemical sensitivity tests, LFLA-215⍰ exhibited resistance to a wide range of antibiotics and compounds, including amikacin, bleomycin, capreomycin, neomycin, gentamicin, kanamycin, polymyxin B, paromomycin, vancomycin, DL-serine hydroxamate, sisomicin, and sulfamethazine, among others (**Table S6**). Susceptibility was observed to chlortetracycline, amoxicillin, cloxacillin, lomefloxacin, minocycline, demeclocycline, cephalothin, ofloxacin, penicillin G, fusidic acid, and others (**Table S6**). Weak growth was observed in the presence of erythromycin, tetracycline, 5-fluoroorotic acid, spiramycin, dodecyltrimethyl ammonium bromide, and related compounds (**Table S6**).

Key phenotypic features distinguishing *Pectobacterium sinaloense* LFLA-215^T^ from other closely related *Pectobacterium* species were identified (Table 2). All compared strains showed growth at 1% NaCl, pH 6, D-saccharic acid, mucic acid, D-galacturonic acid, α-D-glucose, D-mannose, D-fructose, D-galactose, gentiobiose, sucrose, α-D-lactose, β-methyl-D-glucoside, D-salicin, N-acetyl-D-glucosamine, myo-Inositol, D-mannitol, glycerol, D-glucose-6-PO4, D-fructose-6-PO4, D-aspartic acid, L-aspartic acid, L-serine and pectin. Conversely, no growth was observed with propionic acid and D-serine.

**Table 2.** Phenotypic characteristics that differentiate Pectobacterium sinaloense LFLA-215^T^ from closely related Pectobacterium species. Comparative data were obtained from Zhou et al. (2022) for P. colocasium LJ1^T^ and P. aroidearum LJ2, and from Hong et al. (2023) for P. jejuense 13-115 ^T^, P. brasiliense CFBP 6617, P. carotovorum CFBP 2046, P. polare CFBP 8603, and P. parvum CFBP 8630.

| Characteristics | *P. sinaloense* sp.<br>nov. LFLA-215<sup>T</sup> | *P. colocasium*<br>LJ1<sup>T</sup> | *P. aroidearum*<br>LJ2 | *P. jejuense*<br>13-115<sup>T</sup> | *P. brasiliense*<br>CFBP 6617 | *P.*<br>*carotovorum*<br>CFBP 2046 | *P. polare*<br>CFBP 8603 | *P. parvum*<br>CFBP 8630 |
| --- | --- | --- | --- | --- | --- | --- | --- | --- |
| Growth at |  |  |  |  |  |  |  |  |
| pH 5 | + | W | W | - | + | - | - | - |
| 4% NaCl | - | W | W | W | + | - | + | + |
| 8% NaCl | - | - | - | - | W | - | W | - |
| **Carbon source utilization** |  |  |  |  |  |  |  |  |
| Dextrin | + | - | - | - | - | W | - | - |
| D-Maltose | + | - | - | W | - | - | - | - |
| D-Trehalose | + | W | W | + | + | + | + | + |
| D-Cellobiose | + | W | - | + | + | + | + | + |
| D-Turanose | + | - | - | W | + | - | - | - |
| Stachyose | + | - | - | W | + | + | + | + |
| D-Raffinose | W | W | - | + | + | + | + | W |
| D-Melibiose | + | W | - | + | + | + | + | + |
| N-Acetyl-β-D-mannosamine | + | W | W | W | - | - | - | - |
| N-Acetyl-D-galactosamine | W | - | - | - | - | - | - | - |
| N-Acetyl neuraminic acid | W | - | - | - | - | - | - | - |
| 3-Methyl glucose | + | - | W | - | - | - | - | - |
| D-Fucose | + | - | - | - | - | - | - | - |
| L-Rhamnose | + | W | + | + | + | + | + | + |
| L-Galactonic acid lactone | W | + | + | + | - | + | + | + |
| D-Gluconic acid | + | W | W | + | + | + | + | + |
| D-Glucuronic acid | + | W | W | - | - | + | - | W |
| Glucuronamide | + | W | W | - | W | W | - | - |
| Methyl pyruvate | + | + | W | + | - | + | + | + |
| L-Lactic acid | + | - | W | - | + | - | - | - |
| Citric acid | + | - | + | + | + | + | + | - |
| L-Malic acid | + | + | + | + | + | + | + | + |
| Bromo-succinic acid | + | + | + | + | - | + | + | + |
| Tween 40 | + | W | - | W | - | W | - | - |
| α-Keto-butyric acid | W | - | - | - | - | W | - | - |
| Acetoacetic acid | + | W | W | + | W | + | - | - |
| Formic acid | + | W | W | + | - | + | + | + |
| Acetic acid | + | W | + | + | + | + | + | + |
| Inosine | W | - | W | - | - | + | + | + |
| D-Sorbitol | + | - | - | + | - | + | - | - |
| D-Arabitol | + | - | W | + | - | - | - | - |
| L-Alanine | + | - | W | W | - | - | - | - |
| L-Arginine | + | - | - | - | - | - | - | - |
| L-Glutamic acid | W | W | W | + | - | + | + | W |
| **Chemical sensibility** |  |  |  |  |  |  |  |  |
| Aztreonam | - | N/A | N/A | - | + | - | - | - |
| Lincomycin | W | N/A | N/A | - | + | - | - | - |
| Minocycline | - | N/A | N/A | - | + | - | - | - |
| Nalidixic acid | - | N/A | N/A | - | + | - | - | + |
| Potassium tellurite | - | N/A | N/A | - | + | - | + | - |

+ positive reaction, - negative reaction, W weak positive reaction.

The pathogenicity of *Pectobacterium sinaloense* LFLA-215^T^ was confirmed on potato plants (**Figure S3A-E**). Inoculated stems and nearby aerial parts (petioles) exhibited blackening and decay (**Figure S3C-E**), though symptoms only appeared after 6 days post-inoculation, significantly later than those caused by *Pectobacterium versatile* LFLOG-78, where symptoms were visible within 72 h post-inoculation (**Figure S3B**). Similarly, in potato tuber maceration assays, *P. sinaloense* LFLA-215^T^ demonstrated reduced tissue degradation capacity, macerating an average of 2.52 g of potato tuber after 72 h, compared to 6.0 g by *P. versatile* (**Figure S3F**). The delayed symptom development (6 days vs. 72 h) and lower maceration efficiency suggest that while LFLA-215^T^ is pathogenic, it exhibits attenuated virulence compared to *P. versatile* LFLOG-78, a member of the *P. carotovorum* complex. This phenotypic disparity may correlate with LFLA-215 ^T^‘s reduced genome, which ranks at the 8th percentile for size (4.52 Mbp) and 11th percentile for gene count among all *Pectobacterium* genomes with complete chromosome assemblies (**Figure S4**). The genome’s compact architecture, lacking approximately 200–300 genes present in more virulent relatives, could explain its delayed symptom progression and reduced aggressiveness.

## FUNCTIONAL GENOMIC ANALYSES OF STRAIN LFLA-215^T^

Functional annotation of strain LFLA-215^T^ genes was performed using BlastKOALA (Kanehisa et al., 2016), which assigns KEGG Orthology (KO) identifiers through BLAST searches against the non-redundant KEGG database (www.kegg.jp). This enabled the reconstruction of KEGG pathways, BRITE hierarchies, and modules, facilitating inference of high-level biological functions.

Out of all annotated genes, 2,702 (66.8%) were assigned KO identifiers, allowing the reconstruction of 95 complete KEGG pathway modules. These spanned major functional categories such as carbohydrate, nitrogen, lipid, nucleotide, amino acid, and sulfur metabolism, as well as carbon fixation, ATP synthesis, and the biosynthesis of terpenoids and polyketides (**Table S7**). The most represented category was metabolism, particularly carbohydrate metabolism with 325 genes. This was followed by environmental information processing, with a strong presence in membrane transport (251 genes), including ABC transporters, the phosphotransferase system (PTS), and secretion systems. The signal transduction subcategory also showed notable representation (167 genes). Another relevant category was cellular processes, where many genes were linked to virulence-related subcategories, especially cellular community (127 genes), involved in *quorum sensing* and biofilm formation, and cell motility (64 genes), related to chemotaxis and flagellar assembly, consistent with the observed peritrichous flagella and motility of the strain.

A functional genome comparison was conducted between *P. sinaloense* LFLA-215^T^ and all other *Pectobacterium* type strains. Predicted proteins from each strain were annotated with KO identifiers as mentioned above, and functional profiles were compared using KEGG-decoder (Graham et al. 2018). This analysis generated a completeness matrix of predicted metabolic functions, including full pathways, multi-subunit complexes, and individual proteins, allowing comparisons of functional capacity in relation to phylogenetic structure across the genus.

The most notable differences were observed in secretion system completeness (**Figure S5**), confirming the absence of a type IV secretion system in *P. sinaloense* LFLA-215⍰. Another key distinction was found in the CP-lyase operon, where only 12 out of 23 *Pectobacterium* species displayed this predicted function. The CP-lyase complex, encoded by the *phn* operon, is responsible for phosphonate utilization, a potential ecological advantage in phosphorus-limited environments (Adams et al., 2008). Overall, these distinct functional traits further differentiate *P. sinaloense* apart from its closest phylogenetic relatives, *P. aroidearum* and *P. colocasium* (**Figure S6**).

## DESCRIPTION OF PECTOBACTERIUM SINALOENSE SP. NOV

*Pectobacterium sinaloense* etymology: si.na.lo.en’se. N.L. neut. adj. *sinaloense*, pertaining to Sinaloa, the state in Mexico where the type strain LFLA-215^T^ was isolated.

Cells are Gram-negative, motile rods measuring 4.3⍰±⍰0.91µm in length and 1.31⍰±⍰0.09⍰µm in width with peritrichous flagella. The novel taxon is catalase-positive and oxidase-negative. Growth occurs in LB broth at 28⍰°C and 37⍰°C, in the presence of 1% NaCl, and at pH values ranging from 4 to 8. No growth is observed at 4% or 8% NaCl. After 24⍰h of incubation on LB agar at 28⍰°C, colonies are small (1–2⍰mm in diameter), circular, opaque, white-beige with entire margins, convex elevation, moist surfaces, and no diffusible pigment production. LFLA-215^T^ also grows on nutrient agar (NA), crystal violet pectate agar and B-King medium.

BIOLOG phenotypic profiling showed positive utilization of a broad range of carbon sources, including L-arabinose, N-acetyl-D-glucosamine, D-galactose, D-mannose, glycerol, L-proline, sucrose, and many others. Weakly positive reactions were observed with α-ketobutyric acid, 2-hydroxybenzoic acid, itaconic acid, L-methionine, and 2,3-butanedione. LFLA-215^T^ was negative for utilization of propionic acid, capric acid, and D-serine.

The type strain is LFLA-215^T^, isolated in January-2020 from symptomatic potato stems in Ahome, Sinaloa, Mexico (25.8656 N, 108.9249 W). The 16S rRNA sequence of strain LFLA-215^T^ has been deposited in NCBI GenBank and assigned accession number: PV590475.1. The associated BioProject, and BioSample numbers are PRJNA1257438 and SAMN48267889, respectively.

## AKNOWLEDGEMENTS

José Luis Valdez López gratefully acknowledges the financial support provided by SECIHTI (formerly CONACYT; No. 842675), Instituto Politécnico Nacional (IPN), Dirección de Relaciones Internacionales (DRI-IPN), and the PIFI-BEIFI program. RCL is funded by CIHR, Genome Quebec and Genome Canada.

