## Supplementary Material for "*Pectobacterium sinaloense* sp. nov., a novel phytopathogenic species isolated from potato plants in Mexico"

**Table S1.** Tools and databases used by the Mettannotator annotation pipeline v1.3.

| **Tool/Database** | **Version** | **Purpose** | **Reference** |
| --- | --- | --- | --- |
| Bakta | 1.9.3 | CDS calling and functional annotation | (Schwengers et al., 2021) |
| Bakta db | 2024-01-19 | Bakta DB (when Bakta is used as the gene caller) | (Schwengers et al., 2021) |
| InterProScan | 5.62-94.0 | Protein annotation (InterPro, Pfam) | (Jones et al., 2014 |
| eggNOG-mapper | 2.1.11 | Protein annotation (eggNOG, KEGG, COG, GO-terms) | (Cantalapiedra et al., 2021) |
| eggNOG DB | 5.0.2 | Database for eggNOG-mapper | (Cantalapiedra et al., 2021) |
| UniFIRE | 2023.4 | Protein annotation | (MacDougall et al., 2020) |
| AMRFinderPlus | 3.12.8 | Antimicrobial resistance gene annotation; virulence factors, biocide, heat, acid, and metal resistance gene annotation | (Feldgarden et al., 2021) |
| AMRFinderPlus DB | 3.12 2024-01-31.1 | Database for AMRFinderPlus | (Feldgarden et al., 2021) |
| DefenseFinder | 1.2.0 | Annotation of anti-phage systems | (Tesson et al., 2022) |
| DefenseFinder models | 1.2.3 | Database for DefenseFinder | (Tesson et al., 2022) |
| GECCO | 0.9.8 | Biosynthetic gene cluster annotation | (Carroll et al., 2021) |
| antiSMASH | 7.1.0 | Biosynthetic gene cluster annotation | (Blin et al., 2023) |
| SanntiS | 0.9.3.3 | Biosynthetic gene cluster annotation | (Sanchez et al., 2023) |
| run_dbCAN | 4.1.2 | PUL prediction | (Zheng et al., 2023) |
| dbCAN DB | 4.1.3_V12 | Database for run_dbCAN | (Zheng et al., 2023) |
| CRISPRCasFinder | 4.3.2 | Annotation of CRISPR arrays | (Couvin et al., 2018) |
| Rfam | 14.9 | Identification of SSU/LSU rRNA and other ncRNAs | (Kalvari et al., 2018) |
| tRNAscan-SE | 2.0.9 | tRNA predictions | (Chan et al., 2021) |
| Infernal | 1.1 | RNA searches | (Nawrocki & Sean, 2013) |
| pyCirclize | 1.4.0 | Visualize the merged GFF file | (Shimoyama, Y., 2022) |
| Docker | 27.3.1 | Automates the packaging and deployment of software applications using containers. | (Merkel, D., 2014) |
| Nextflow | 24.04.4 | Workflow management system with container support used to run the pipeline ebi-metagenomics/mettannotator: '1.3'. | (Di Tommaso et al., 2017) |

**Table S2.** Genome assembly and annotation metrics of *Pectobacterium sinaloense* LFLA-215^T^.

| **Characteristic** | | ***Pectobacterium sinaloense* LFLA-215^T^** |
| --- | --- | --- |
| *De novo* assembly | Accession number | - |
|  | Genome size (bp) | 4,522,015 |
|  | GC ratio | 0.515826683 |
|  | # Ambiguous bases | 0 |
|  | # Contigs | 1 |
|  | Largest contig (bp) | 4,522,015 |
|  | N50 (bp) | 4,522,015 |
|  | L50 | 1 |
|  | Coverage | 632x |
| CheckM analysis | Completeness (%) | 93.33 |
|  | Contamination (%) | 0.44 |
| Annotation | Coding sequences | 4,043 |
|  | Coding density | 0.862435441 |
|  | # BGC ( identified by antiSMASH) | 6 |
|  | # BGC (Identified by Gecco) | 4 |
|  | # BGC (Identified by SanntiS) | 10 |
|  | # PUL (Identified by dbcan3) | 63 |
|  | AMR genes (Identified by AMRFinderPlus) | 3 |
|  | Antiphage-defense systems (genes) | 11 (38) |
|  | tRNAs | 77 |
|  | Hypothetical proteins | 164 |
|  | PCWDEs genes | 33 |
|  | Secretion systems (genes) | 6 (48) |
|  | Toxin-related genes | 34 |

BGC; Biosynthetic gene cluster, PUL; Polysaccharide utilization loci, AMR; Antimicrobial resistance, PCWDEs; Plant cell wall-degrading enzymes.

**Table S3.** Key genomic regions in *Pectobacterium sinaloense* LFLA-215^T^ identified by AntiSMASH, Gecco, SanntiS, DefenseFinder and dbCAN3.

| **Region** | **Predicted Substrate/Synthesis/Function** | **From** | **To** | **Tool** |
| --- | --- | --- | --- | --- |
| PUL | N/A | 131558 | 135509 | dbCAN3 |
| PUL | arabinan | 196499 | 201422 | dbCAN3 |
| BGC | betalactone | 202842 | 226656 | AntiSMASH |
| PUL | N/A | 228285 | 232977 | dbCAN3 |
| PUL | N/A | 238887 | 243705 | dbCAN3 |
| PUL | N/A | 264911 | 272383 | dbCAN3 |
| PUL | pectin | 302775 | 307058 | dbCAN3 |
| PUL | cellobiose | 340109 | 343853 | dbCAN3 |
| PUL | N/A | 347541 | 353040 | dbCAN3 |
| PUL | pectin | 439189 | 441931 | dbCAN3 |
| PUL | N/A | 447479 | 450683 | dbCAN3 |
| PUL | xylan | 515825 | 522665 | dbCAN3 |
| Anti-phage system | CAS_Class1-Subtype-III-A | 564424 | 574497 | DefenseFinder |
| PUL | N/A | 665334 | 678841 | dbCAN3 |
| PUL | N/A | 705321 | 712154 | dbCAN3 |
| PUL | N/A | 782282 | 790045 | dbCAN3 |
| PUL | pectin | 792785 | 807306 | dbCAN3 |
| PUL | starch | 820659 | 823614 | dbCAN3 |
| PUL | galactan | 872756 | 895177 | dbCAN3 |
| BGC | Saccharide | 922588 | 931048 | SanntiS |
| PUL | N/A | 927178 | 938119 | dbCAN3 |
| BGC | NRP | 937796 | 946477 | SanntiS |
| Anti-phage system | RM_Type_I | 946623 | 952346 | DefenseFinder |
| PUL | N/A | 960282 | 982626 | dbCAN3 |
| BGC | NRP | 1010632 | 1017857 | SanntiS |
| Anti-phage system | SEFIR | 1157890 | 1159326 | DefenseFinder |
| Anti-phage system | Gao_Qat | 1171400 | 1176134 | DefenseFinder |
| Anti-phage system | RM_Type_I | 1176191 | 1182852 | DefenseFinder |
| Anti-phage system | Azaca | 1184710 | 1190868 | DefenseFinder |
| Anti-phage system | Eleos | 1191039 | 1195121 | DefenseFinder |
| PUL | N/A | 1244245 | 1251024 | dbCAN3 |
| PUL | N/A | 1277842 | 1284072 | dbCAN3 |
| PUL | cellobiose | 1391355 | 1397907 | dbCAN3 |
| PUL | galactan | 1478560 | 1496997 | dbCAN3 |
| PUL | N/A | 1502435 | 1511803 | dbCAN3 |
| PUL | N/A | 1619617 | 1620613 | dbCAN3 |
| Anti-phage system | Gao_Tmn | 1700860 | 1704660 | DefenseFinder |
| PUL | pectin | 1766029 | 1785475 | dbCAN3 |
| PUL | cellobiose | 1800199 | 1803162 | dbCAN3 |
| PUL | N/A | 1831086 | 1833817 | dbCAN3 |
| PUL | N/A | 1841065 | 1844802 | dbCAN3 |
| PUL | levoglucosan | 1888153 | 1895069 | dbCAN3 |
| PUL | pectin | 2012660 | 2027898 | dbCAN3 |
| PUL | pectin | 2036234 | 2048519 | dbCAN3 |
| PUL | N/A | 2095070 | 2102343 | dbCAN3 |
| BGC | thiopeptide | 2199698 | 2226123 | AntiSMASH |
| PUL | N/A | 2351530 | 2357015 | dbCAN3 |
| PUL | xylan | 2364246 | 2370769 | dbCAN3 |
| PUL | N/A | 2386102 | 2391959 | dbCAN3 |
| PUL | pectin | 2507205 | 2556037 | dbCAN3 |
| PUL | host glycan | 2560098 | 2565927 | dbCAN3 |
| PUL | levoglucosan | 2570520 | 2575290 | dbCAN3 |
| PUL | N/A | 2582578 | 2590846 | dbCAN3 |
| BGC | Saccharide | 2584796 | 2615963 | Gecco |
| BGC | Saccharide | 2586515 | 2613986 | SanntiS |
| PUL | capsule polysaccharide synthesis | 2599491 | 2615963 | dbCAN3 |
| PUL | levoglucosan | 2640745 | 2651943 | dbCAN3 |
| PUL | glycosaminoglycan | 2719797 | 2723608 | dbCAN3 |
| PUL | N/A | 2830083 | 2841763 | dbCAN3 |
| PUL | N/A | 2931801 | 2937013 | dbCAN3 |
| PUL | N/A | 2943071 | 2947110 | dbCAN3 |
| PUL | N/A | 2996868 | 3005872 | dbCAN3 |
| PUL | beta-glucoside | 3117335 | 3137123 | dbCAN3 |
| Anti-phage system | AbiE | 3158236 | 3159682 | DefenseFinder |
| PUL | xylan | 3162888 | 3168496 | dbCAN3 |
| PUL | N/A | 3184516 | 3197328 | dbCAN3 |
| PUL | N/A | 3226623 | 3234769 | dbCAN3 |
| PUL | melibiose | 3274523 | 3287290 | dbCAN3 |
| BGC | RiPP | 3306932 | 3312352 | SanntiS |
| BGC | NRPS,T1PKS | 3317326 | 3373474 | AntiSMASH |
| BGC | NRP Polyketide | 3333440 | 3357331 | SanntiS |
| BGC | NRP;Polyketide | 3334484 | 3355234 | Gecco |
| PUL | cellobiose | 3380031 | 3385678 | dbCAN3 |
| BGC | NRP-metallophore,NRPS | 3409416 | 3463413 | AntiSMASH |
| BGC | NRP | 3421558 | 3450347 | SanntiS |
| BGC | NRP | 3427317 | 3450347 | Gecco |
| Anti-phage system | CAS_Class1-Subtype-I-E | 3539701 | 3549598 | DefenseFinder |
| PUL | starch | 3551990 | 3564981 | dbCAN3 |
| BGC | Saccharide | 3785881 | 3805937 | SanntiS |
| PUL | N/A | 3788276 | 3795868 | dbCAN3 |
| Anti-phage system | RM_Type_IV | 3850738 | 3852576 | DefenseFinder |
| BGC | Homoserine lactone | 3864628 | 3885281 | AntiSMASH |
| BGC | Unknown | 3868165 | 3879146 | Gecco |
| PUL | pectin | 3870420 | 3873526 | dbCAN3 |
| PUL | N/A | 4017317 | 4026773 | dbCAN3 |
| PUL | host glycan | 4068200 | 4086483 | dbCAN3 |
| PUL | chitin | 4091242 | 4103129 | dbCAN3 |
| PUL | cellulose | 4131794 | 4148203 | dbCAN3 |
| BGC | Saccharide | 4308468 | 4324922 | SanntiS |
| PUL | N/A | 4316171 | 4320637 | dbCAN3 |
| PUL | glycogen | 4363756 | 4388715 | dbCAN3 |
| BGC | NI-siderophore | 4399991 | 4429892 | AntiSMASH |
| BGC | Other | 4412301 | 4427199 | SanntiS |
| PUL | pectin | 4459427 | 4470277 | dbCAN3 |

*PUL; polysaccharide utilization loci and BGC; biosynthetic gene clusters, N/A; Not Available

**Table S4.** Genes identified in *Pectobacterium sinaloense* LFLA-215^T^ grouped by functional category, including plant cell wall-degrading enzymes (PCWDEs), secretion system components (types I, II, III, and VI), and toxin-related genes.

| **Category** | | **Genes** |
| --- | --- | --- |
| PCWDEs | Pectinase | *YesR_1, YesR_2, Pgl, PemA, PaeY, Pel1, Pgu1, FhaB,* Pectate lyase *PlyH/PlyE-*like*, PelB, KdgF, PelW,* Oligogalacturonide lyase*, PelN,* alpha-L-rhamnosidase*, pnl, Pgu1_2, pehA, PelI,* Rhamnogalacturonan lyase*, PemB, PelX, FhaB, PelZ, Pel3, Pel2,* Beta-xylosidase |
|  | Cellulase | *BcsZ, celV1* |
|  | Protease | *Prt1,* Serralysin*, pqqL,* Acetulornithine deacetylase |
| Secretion secretions | T1SS (3) | *(EmrA, sunT* and *TolC), (TolC, EmrA* and *arpD) and (*ABC transporter type *1, MFP, TolC)* |
|  | T2SS | *GspC, GspD, GspE, GspF, GspG, GspH,GspI, GspJ, GspK, GspL, GspM, GspN, GspO* |
|  | T3SS | *SctU, SctT, SctS, SctR, SctQ, HrpP,* Type III secretion protein *(I), SctN, SctD, SctV, SctW, HrpB, SctJ,* Type III secretion protein *(II), SctL, HrpF,* Type III secretion protein *(III), SctC, HrpT,* Type III secretion protein *(III)* |
|  | T6SS | *TssB, TssC, TssE, TssF, TssG, TssJ, TssK, DotU, TssH, TssA, TssM, EvfE, VgrGB, Zn-binding (PAAR), Hcp, vgrG, evpJ* |
| Toxins | | *Tab, Hcp (I), SymE (I), SymE (II),* Bacterial toxin 50 domain-containing protein*, SymE (III), Nip, CvpA, CbtA, OrtT,* VENN motif containing protein *(I),* VENN motif containing protein *(II), tnt,* VENN motif containing protein *(III), HigB2 (I), AvrE, HrpK, SrfC, ParE, HicA, PasT, AbiEii, RelE/ParE-like (I), CptA, Hcp (II), YefM, YoeB, RhaS, RelE/ParE-like (II), RelE (I), SymE (IV), Hcp (III), RelE (II), HigB2 (II)* |

**Table S5.** Bacterial dimension measurements using transmission electron microscopy.

| **Measured cell** | **Length (µm)** | **Width (µm)** |
| --- | --- | --- |
| 1 | 2.99 | 1.227 |
| 2 | 4.313 | 1.092 |
| 3 | 4.098 | 1.213 |
| 4 | 4.702 | 1.349 |
| 5 | 2.987 | 1.314 |
| 6 | 5.299 | 1.217 |
| 7 | 5.252 | 1.427 |
| 8 | 4.235 | 1.202 |
| 9 | 4.731 | 1.333 |
| 10 | 4.603 | 1.453 |
| 11 | 4.717 | 1.266 |
| 12 | 3.213 | 1.392 |
| 13 | 5.183 | 1.414 |
| 14 | 6.236 | 1.399 |
| 15 | 3.042 | 1.354 |
| Average | 4.3734 | 1.310133333 |
| Stand. Desv. | 0.908767696 | 0.096718578 |

**Table S6.** Complete results of carbon source utilization and chemical sensitivity tests for *Pectobacterium sinaloense* LFLA-215^T^, based on BIOLOG phenotypic assays. The table includes positive, negative, and positive weak reactions to various carbon sources and chemical compounds.

| **Biochemical test** | **Results** | **Compounds** |
| --- | --- | --- |
| **Carbon source utilization** | **Positive** | L-Arabinose, N-Acetyl-D-Glucosamine, D-Saccharic acid, Succinic acid, D-Galactose, L-Aspartic acid, L-Proline, D-Alanine, D-Trehalose, D-Mannose, Dulcitol, D-Sorbitol, Glycerol, L-Fucose, D-Glucuronic acid, D-Gluconic acid, DL-a-Glycerol Phosphate, D-Xylose, L-Lactic acid, Formic acid, D-Mannitol, L-Glutamic acid, D-Glucose-6-Phosphate, D-Galactonic acid-g-Lactone, DL-Malic acid, D-Ribose, Tween 20, L-Rhamnose, D-Fructose, Acetic acid, a-D-Glucose, Maltose, D-Melibiose, Thymidine, L-Asparagine, D-Aspartic acid, D-Glucosaminic acid, 1,2-Propanediol, Tween 40, a-Ketoglutaric acid, a-Methyl-D-Galactoside, a-D-Lactose, Lactulose, Sucrose, Uridine, L-Glutamine, m-Tartaric acid, D-Glucose-1-Phosphate, D-Fructose-6-Phosphate, Tween 80, a-Hydroxyglutaric acid-g-Lactone, a-Hydroxybutyric acid, b-Methyl-D-Glucoside, Adonitol, Maltotriose, 2`-Deoxyadenosine, Adenosine, Gly-Asp, Citric acid, m-Inositol, D-Threonine, Fumaric acid, Bromosuccinic acid, Mucic acid, Glycolic acid, Glyoxylic acid, D-Cellobiose, Inosine, Gly-Glu, Tricarballylic acid, L-Serine, L-Threonine, L-Alanine, Ala-Gly, Acetoacetic acid, N-Acetyl-D-Mannosamine, Mono-Methylsuccinate, Methylpyruvate, D-Malic acid, L-Malic acid, Gly-Pro, p-Hydroxyphenyl Acetic acid, m-Hydroxyphenyl Acetic acid, Tyramine, D-Psicose, L-Lyxose, Glucuronamide, Pyruvic acid, L-Galactonic acid-g-Lactone, D-Galacturonic acid, Phenylethylamine, 2-Aminoethanol, Chondroitin Sulfate C, a-Cyclodextrin, b-Cyclodextrin, g-Cyclodextrin, Dextrin, Gelatin, Glycogen, Inulin, Laminarin, Mannan, Pectin, N-Acetyl-D-Galactosamine, N-Acetyl-Neuraminic acid, b-D-Allose, Amygdalin, D-Arabinose, D-Arabitol, L-Arabitol, Arbutin, 2-Deoxy-D-Ribose, i-Erythritol, D-Fucose, 3-O-b-D-Galactopyranosyl-D-Arabinose, Gentiobiose, L-Glucose, D-Lactitol, D-Melezitose, Maltitol, a-Methyl-D-Glucoside, b-Methyl-D-Galactoside, 3-Methylglucose, b-Methyl-D-Glucuronic acid, a-Methyl-D-Mannoside, b-Methyl-D-Xyloside, Palatinose, D-Raffinose, Salicin, Sedoheptulosan, L-Sorbose, Stachyose, D-Tagatose, Turanose, Xylitol, N-Acetyl-D-Glucosaminitol, g-Amino Butyric acid, d-Amino Valeric acid, Butyric acid, Hexanoic acid, Citraconic acid, Citramalic acid, D-Glucosamine, 4-Hydroxybenzoic acid, b-Hydroxybutyric acid, g-Hydroxybutyric acid, a-Keto-Valeric acid, 5-Keto-D-Gluconic acid, D-Lactic acid Methyl Ester, Malonic acid, Melibionic acid, Oxalic acid, Oxalomalic acid, Quinic acid, D-Ribono-1,4-Lactone, Sebacic acid, Sorbic acid, Succinamic acid, D-Tartaric acid, L-Tartaric acid, Acetamide, L-Alaninamide, N-Acetyl-L-Glutamic acid, L-Arginine, Glycine, L-Histidine, L-Homoserine, Hydroxy-L-Proline, L-Isoleucine, L-Leucine, L-Lysine, L-Ornithine, L-Phenylalanine, L-Pyroglutamic acid, L-Valine, DL-Carnitine, sec-Butylamine, DL-Octopamine, Putrescine, Dihydroxyacetone and 2,3-Butanediol |
|  | **Weak** | a-Ketobutyric acid, 2-Hydroxybenzoic acid, Itaconic acid, L-Methionine and 2-3-Butanedione |
|  | **Negative** | Propionic acid, Capric acid and D-serine |
| **Chemical sensibility** | **Positive** | Amikacin, Bleomycin, Capreomycin, Neomycin, Gentamicin, Kanamycin, Polymyxin B, Paromomycin, Vancomycin, DL-Serine hydroxamate, Sisomicin, Sulfamethazine, 2,4-Diamino-6,7-diisopropylpteridine, Sulfadiazine, Tobramycin, Sulfathiazole, Spectinomycin, Sulfamethoxazole, D-Cycloserine, 5-7-Dichloro-8-hydroxyquinaldine, Domiphen bromide, Nordihydroguaiaretic acid, Alexidine, Menadione- sodium bisulfite, Zinc chloride, Phosphomycin, Sulfanilamide, Cetylpyridinium chloride, Streptomycin, 5-Azacytidine, Rifamycin SV, Sodium Selenite, Aluminum sulfate, Chromium (III) chloride, Ferric chloride, L-Glutamic acid g-monohydroxamate, b-Chloro-L-alanine, Thiosalicylate, Sulfachloropyridazine, Sulfamonomethoxine, Compound 48/80, Tannic acid, Chlorambucil, Sodium pyrophosphate, Trifluorothymidine, Azathioprine, Poly-L-lysine, Sulfisoxazole, Sodium Bromate, Sodium metasilicate, Triclosan, 3,5- Diamino-1,2,4-triazole (Guanazole), Myricetin, 5-Fluoro-5`-deoxyuridine, 2- Phenylphenol, Plumbagin, Apramycin, Benserazide, Proflavine, Crystal violet and Dodine (n-Dodecylguanidine) |
|  | **Weak** | Erythromycin, Tetracycline, 5-Fluoroorotic acid, Spiramycin, Dodecyltrimethyl ammonium bromide, Cefmetazole, EDTA, 5,7-Dichloro-8-hydroxyquinoline, 5-Nitro-2-furaldehyde semicarbazone (Nitrofurazone), Methyl viologen, Oleandomycin- phosphate salt, Norfloxacin, Dichlofluanid, Hygromycin B, Ethionamide, 3-Amino-1,2,4-triazole, Niaproof, Cefsulodin, Caffeine, Thiamphenicol, Pentachlorophenol, Sodium Arsenite, Lidocaine, Antimony (III) chloride, Semicarbazide hydrochloride, DL-Propanolol, Thioridazine, 18-Crown-6 ether, Hexachlorophene, Pridinol and 8-Hydroxyquinoline |
|  | **Negative** | Chlortetracycline, Amoxicillin, Cloxacillin, Lomefloxacin, Minocycline, Demeclocycline, Cephalothin, Ofloxacin, Penicillin G, Fusidic acid- sodium salt, Phleomycin, Carbonyl-cyanide m-chlorophenylhydrazone (CCCP), Sodium Azide, Cefotaxime, 5-Chloro-7-iodo-8-hydroxy-quinoline, DL-Methionine hydroxamate, Tinidazole, Aztreonam, Ornidazole, Oxytetracycline, Captan and Tolylfluanid |

**Table S7.** Number of genes in the *Pectobacterium sinaloense* LFLA-215^T^ genome assigned to each KEGG defined function by using the BlastKOALA tool.

| **Category** | **Subcategory** | **Genes Count** |
| --- | --- | --- |
| **Metabolism** |  |  |
|  | Carbohydrate metabolism | 325 |
|  | Energy metabolism | 156 |
|  | Lipid metabolism | 64 |
|  | Nucleotide metabolism | 99 |
|  | Amino acid metabolism | 226 |
|  | Metabolism of other amino acids | 57 |
|  | Glycan biosynthesis and metabolism | 113 |
|  | Metabolism of cofactors and vitamins | 183 |
|  | Metabolism of terpenoids and polyketides | 31 |
|  | Biosynthesis of other secondary metabolites | 48 |
|  | Xenobiotics biodegradation and metabolism | 43 |
| **Genetic Information Processing** |  |  |
|  | Transcription | 4 |
|  | Translation | 79 |
|  | Folding, sorting and degradation | 53 |
|  | Replication and repair | 80 |
|  | Information processing in viruses | 1 |
| **Environmental Information Processing** |  |  |
|  | Membrane transport | 251 |
|  | Signal transduction | 167 |
| **Cellular Processes** |  |  |
|  | Transport and catabolism | 7 |
|  | Cell growth and death | 18 |
|  | Cellular community - prokaryotes | 127 |
|  | Cell motility | 64 |
| **Organismal Systems** |  |  |
|  | Immune system | 6 |
|  | Endocrine system | 17 |
|  | Digestive system | 9 |
|  | Nervous system | 2 |
|  | Development and regeneration | 1 |
|  | Aging | 11 |
|  | Environmental adaptation | 7 |
| **Human Diseases** |  |  |
|  | Cancer: overview | 17 |
|  | Cancer: specific types | 4 |
|  | Infectious disease: viral | 1 |
|  | Infectious disease: bacterial | 25 |
|  | Infectious disease: parasitic | 3 |
|  | Immune disease | 2 |
|  | Neurodegenerative disease | 8 |
|  | Cardiovascular disease | 11 |
|  | Endocrine and metabolic disease | 6 |
|  | Drug resistance: antimicrobial | 56 |
|  | Drug resistance: antineoplastic | 6 |

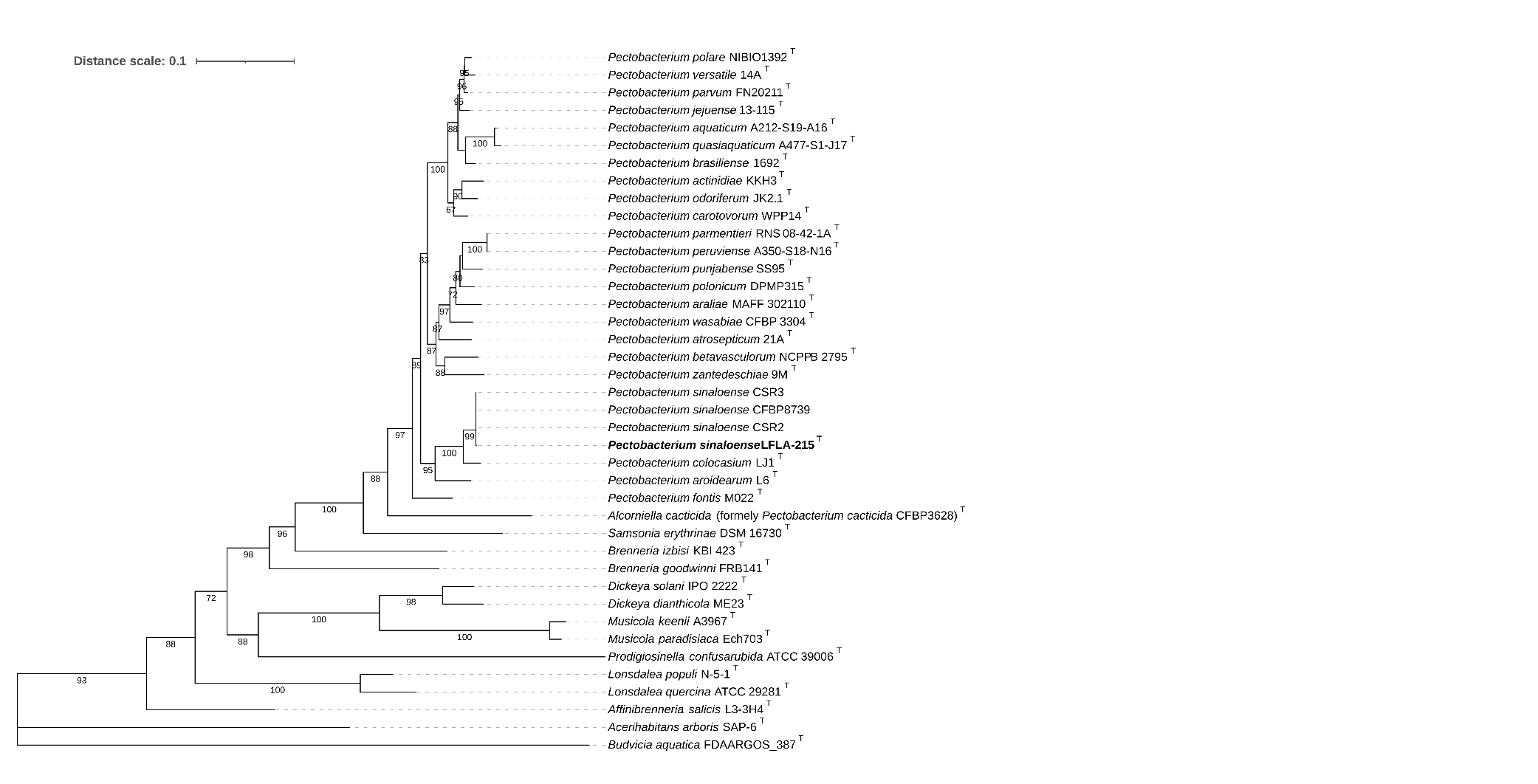
**Figure S1.** Maximum-likelihood phylogenomic tree reconstructed based on the concatenated alignment of the *dnaJ* gene sequences, showing the relationships between *Pectobacterium sinaloense* LFLA-215^T^ (bold type) and closely related strains from the family *Pectobacteriaceae* (listed in Table 1). The total alignment length was 1,143 bp and included 40 sequences, with 45.66% parsimony-informative sites. Phylogenetic inference was performed using IQ-TREE v3.0.0 under the TIM+F+I+G4 model with 1,000 ultrafast bootstrap replicates. Ultrafast bootstrap values are indicated at the nodes (only values ≥60 are shown). *Budvicia aquatica* FDAARGOS_387^T^ was used as the outgroup.

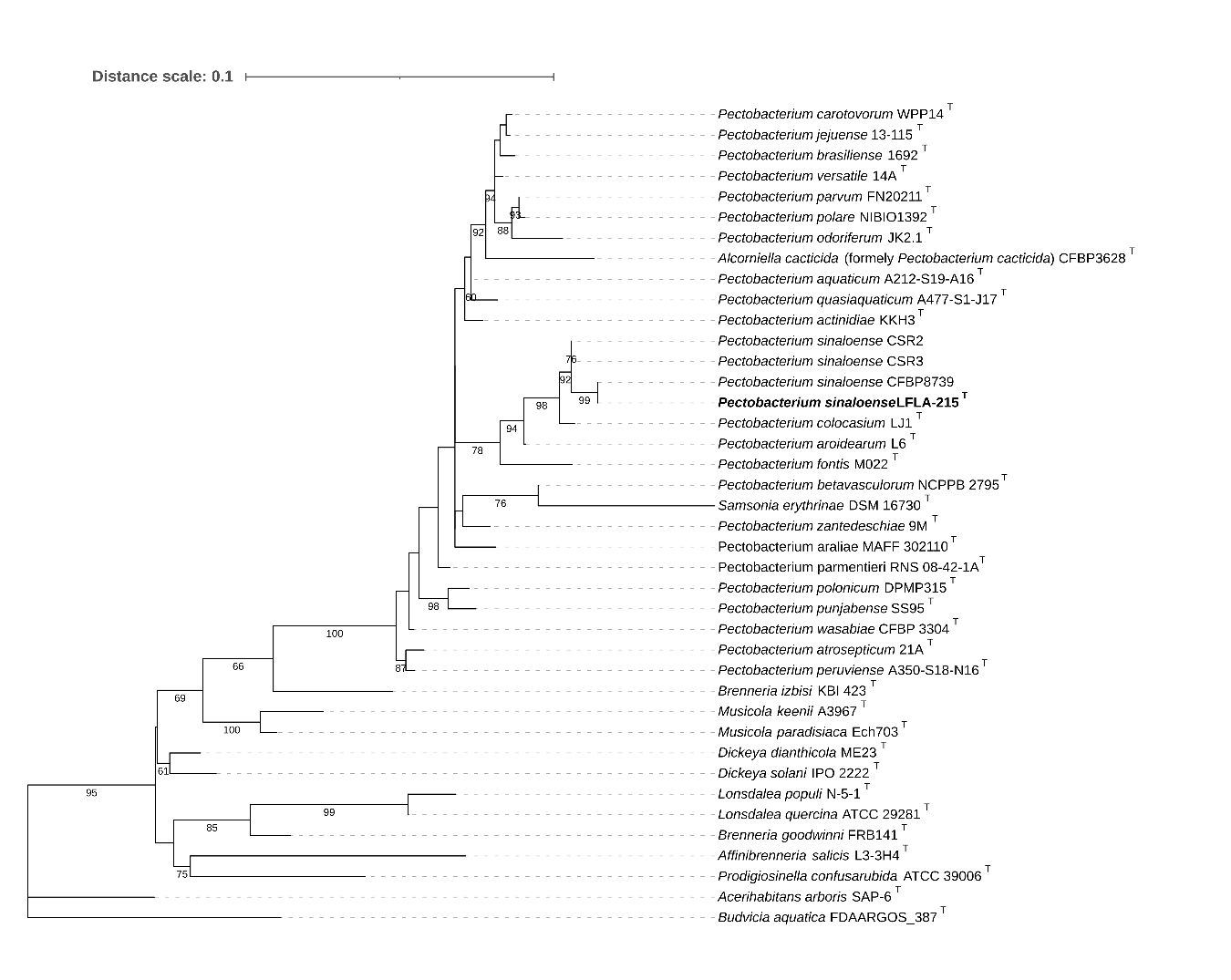
**Figure S2.** Maximum-likelihood phylogenomic tree reconstructed based on the concatenated alignment of 16S ribosomal RNA sequences, showing the relationships between *Pectobacterium sinaloense* LFLA-215^T^ (bold type) and closely related strains from the family *Pectobacteriaceae* (listed in Table 1). The total alignment length was 1,471 bp and included 40 sequences, with 13.59% parsimony-informative sites. Phylogenetic inference was performed using IQ-TREE v3.0.0 under the TPM3u+F+I+G4 model with 1,000 ultrafast bootstrap replicates. Ultrafast bootstrap values are indicated at the nodes (only values ≥60 are shown). *Budvicia aquatica* FDAARGOS_387^T^ was used as the outgroup.

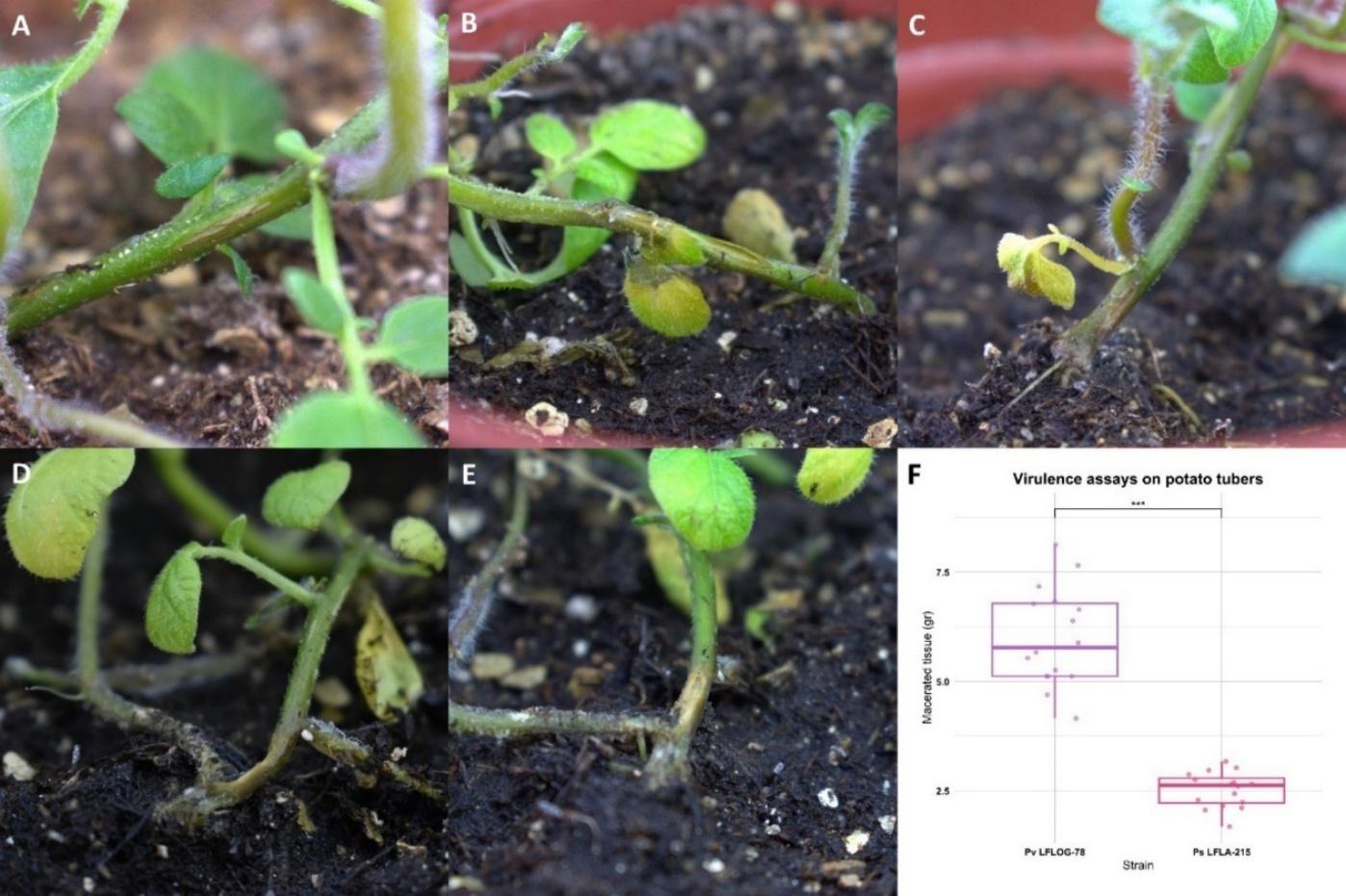

**Figure S3.** Pathogenicity and virulence assays of *Pectobacterium sinaloense* (Ps) LFLA-215^T^ compared to *Pectobacterium versatile* (Pv) LFLOG-78.

**(A)** Mock-inoculated area with sterile 10 mM MgSO_4_; **(B)** symptoms of tissue decay and liquefaction caused by Pv LFLOG-78 three days post-inoculation; **(C–E)** symptoms of tissue decay, blackening, and liquefaction caused by Ps LFLA-215^T^ six days post-inoculation; **(F)** results of virulence assays on potato tubers for Pv LFLOG-78 and Ps LFLA-215^T^ after 72 h. Experiments were performed in duplicate. Statistical analyses were conducted for virulence assay data (n = 16), using the Kruskal–Wallis test followed by Dunn’s post hoc test with Bonferroni correction.

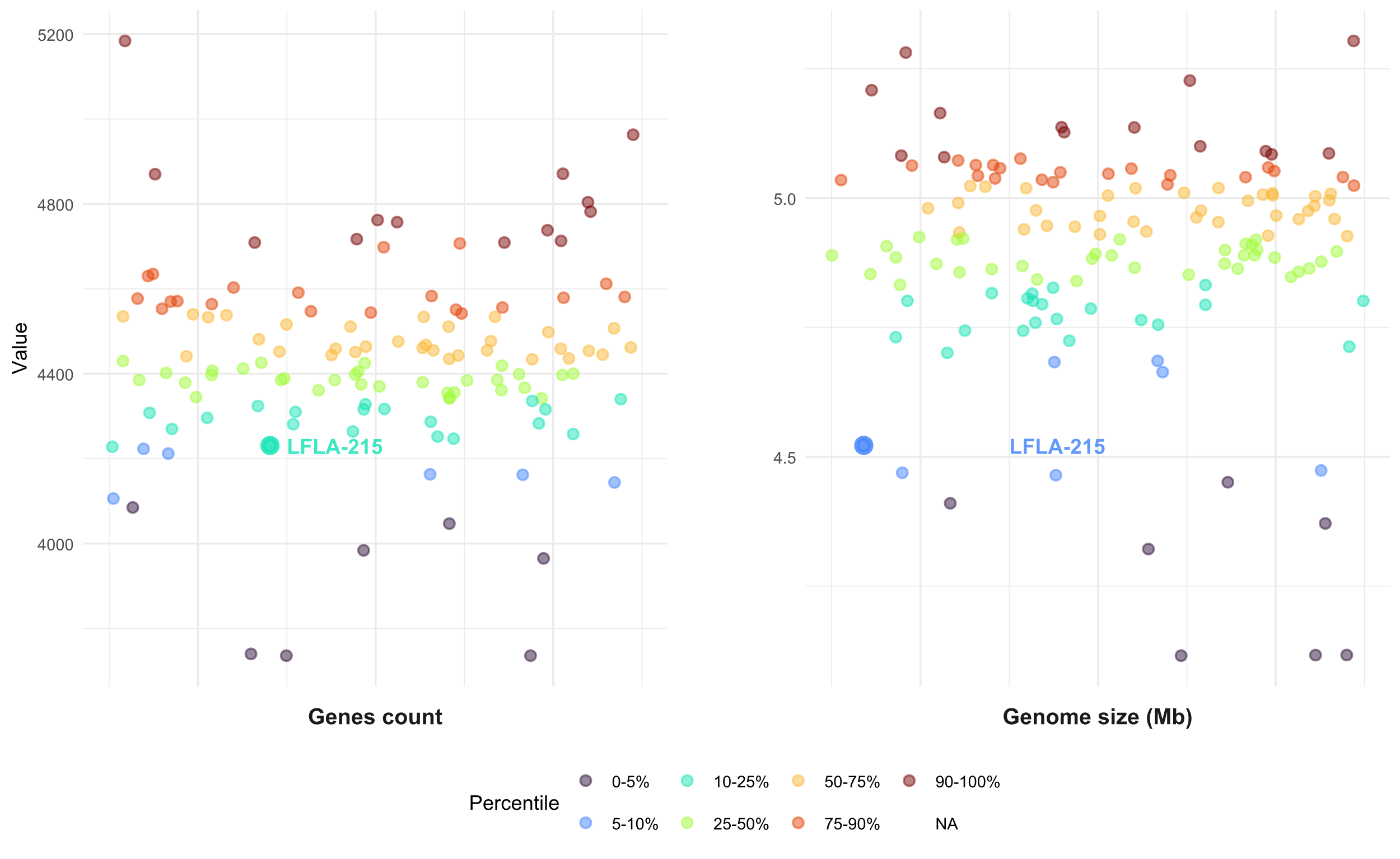
**Figure S4.** Percentile distribution of genome size and gene count in *Pectobacterium* genomes with complete chromosome-level assemblies. LFLA-215^T^ is highlighted (thicker data point) in both distributions.

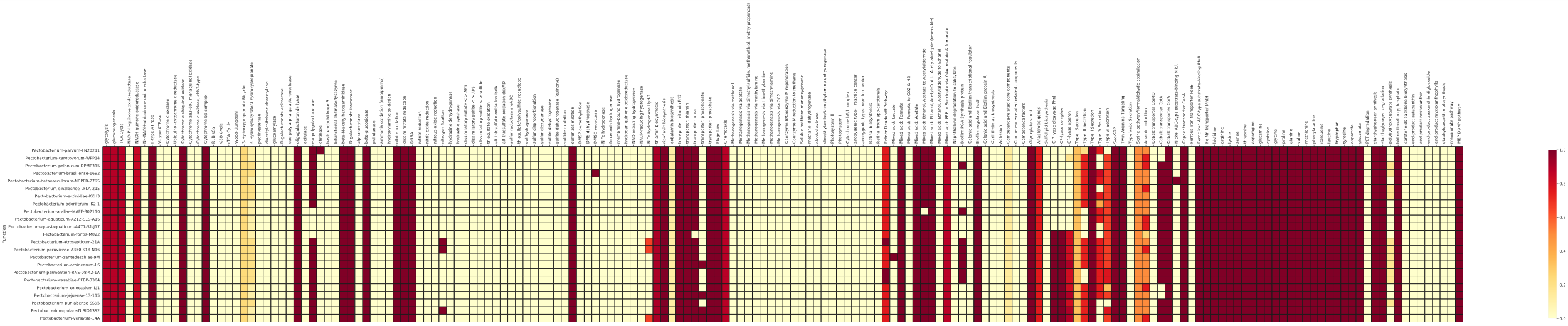

**Figure S5.** Metabolic function completeness matrix for *Pectobacterium sinaloense* LFLA-215^T^ and all *Pectobacterium* type strains. Completeness matrix of predicted metabolic processes identified using KEGG-Decoder, including entire pathways, multi-subunit complexes, and individual proteins

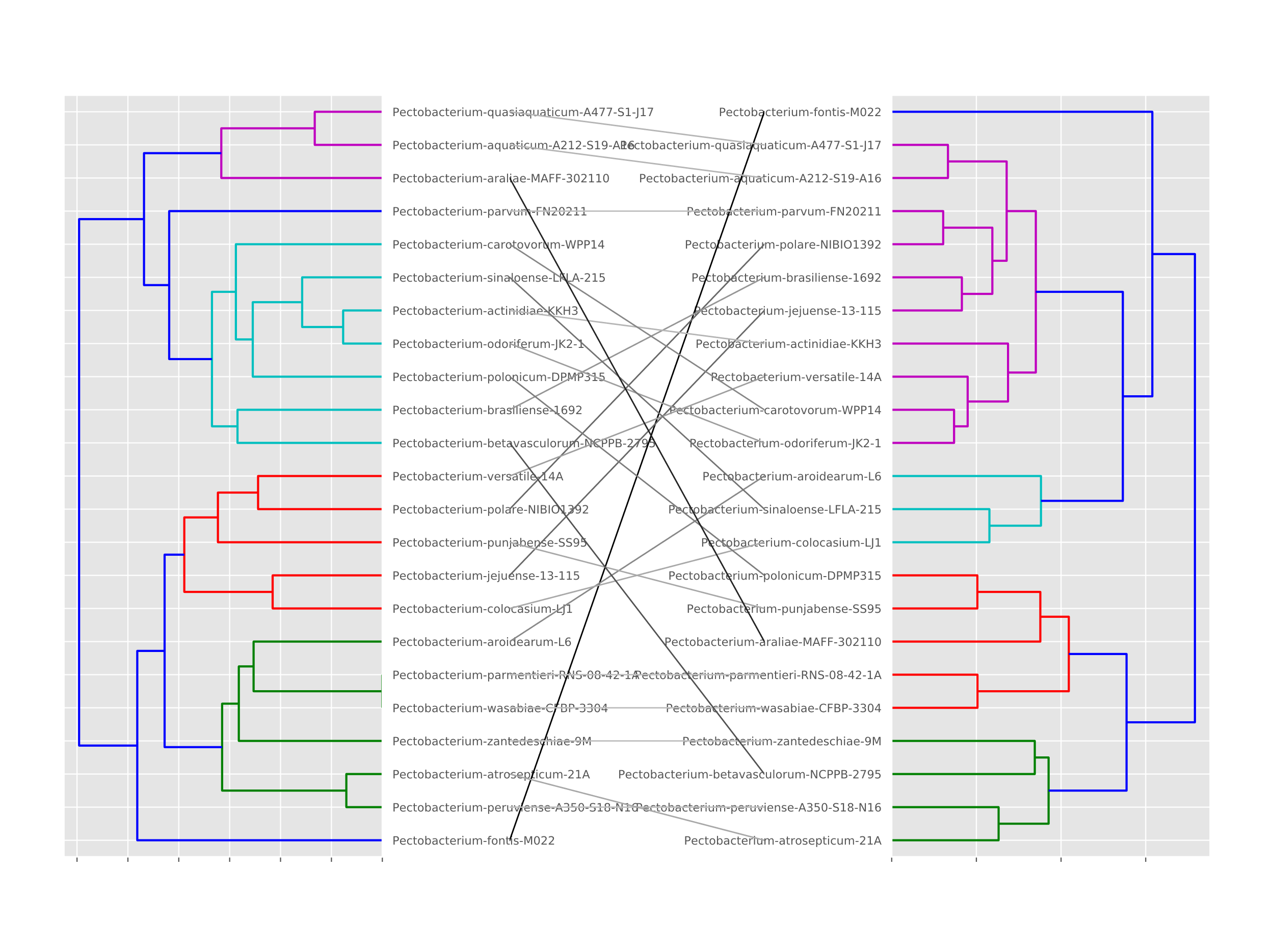

**Figure S6.** Tanglegram illustrating functional and phylogenetic relationships among Pectobacterium type strains. Tanglegram showing the relationship between functional capacities (left dendrogram, based on metabolic profiles from KEGG annotations) and phylogenetic relationships (right dendrogram, inferred from a concatenated alignment of 926 core genes).
